# Glycosylation Limits Forward Trafficking of the Tetraspan Membrane Protein PMP22

**DOI:** 10.1101/2020.12.18.423452

**Authors:** Justin T. Marinko, Madison T. Wright, Darren R. Heintzman, Lars Plate, Charles R. Sanders

## Abstract

Peripheral myelin protein 22 (PMP22) folds and traffics inefficiently, a phenomenon closely related to the mechanisms by which this tetraspan membrane protein causes Charcot-Marie-Tooth disease (CMTD). We report that elimination of N-glycosylation results in a 3-fold increase in the cell surface trafficking of wild type (WT) PMP22 and a 10-fold increase in trafficking of the unstable L16P disease mutant form. Studies of the interactions of PMP22 with oligosaccharyltransferases A and B as well as quantitative proteomic experiments established that critical endoplasmic reticulum (ER) quality control decisions occur earlier in the biogenesis to cell surface trafficking pathway for the L16P mutant than for WT. CRISPR knock-out cell lines for ER proteins calnexin, RER1, and UGGT1 illuminated the role of each protein in glycosylation dependent and independent surface trafficking of WT PMP22, as well as for a series of disease mutants of varying folding stabilities.

**One Sentence Summary:** N-linked glycosylation was seen to dramatically limit the cell surface trafficking of PMP22, with some key quality control factors in PMP22 biogenesis being identified.

## Introduction

Secreted and transmembrane proteins comprise over one third of the human proteome, passing through the endoplasmic reticulum (ER) *en route* to their ultimate destinations^1–3^. For most integral membrane proteins, integration into the ER membrane is intimately coupled to translation. Translating ribosomes associate with the Sec61-translocon complex to thread transmembrane (TM) segments through a central water exposed pore^4–6^. This pore contains a lateral gate that allows translocating polypeptides to sample both lipid and hydrophilic environments^7^. Soluble secretary proteins translocate completely through the pore to enter the ER lumen, while TM helices partition laterally into the ER membrane. Most membrane proteins adopt their secondary structure and attain correct membrane topology at this initial stage of assembly. Once fully synthesized, proteins are released from the translocon, diffuse away, and begin the second stage of membrane protein folding: formation of tertiary and quaternary structure^1, 8^. Once properly folded, proteins traffic beyond the ER via exit sites (ERES) to the Golgi complex and from there to their destination membrane^2^. Proteins that fail to adopt proper structure are retained in the ER to provide additional time for folding or, failing that, are targeted for elimination either by ER-associated degradation (ERAD) or ER-associated autophagy (ER-phagy)^9^.

Protein folding in the ER is under constant surveillance by the resident ER quality control network (ERQC)^1, 10–11^, which contains numerous folding sensors, chaperones, and other proteins, including those involved in ERAD and ER-phagy. Collectively, these proteins monitor, assist, and execute logic decisions as to whether to retain, degrade, or authorize exit of nascent proteins from the ER. Much is known about the molecular details of this pathway for soluble proteins, but less is understood about this process for TM proteins. In this work, we seek to expand our understanding of how ERQC manages quality control decision for human peripheral myelin protein 22 (PMP22).

PMP22 is a tetraspan integral membrane protein (**Figure 1A**) that is highly expressed in the plasma membrane (PM) of myelinating Schwann cells in the peripheral nervous system (PNS)^12–13^. The specific functions of PMP22 are not well-understood^14–16^, but include a structural role in both the maintenance and development of compact myelin^17^. PMP22 shares ~60% sequence similarity with claudin-15, one of the structural proteins involved in maintaining tight junctions^18^. Moreover, when reconstituted into liposomes, PMP22 can induce flattening and wrapping of the vesicles to form myelin-like assemblies^19^. Mutations in the *pmp22* gene, including gene duplication, gene deletion, or any one of more than 40 known single nucleotide polymorphisms, cause a range of progressive peripheral neuropathies including Charcot-Marie-Tooth disease types 1A and E, hereditary neuropathy with liability to pressure palsies (HNPP), and Dejerine-Sottas syndrome (DSS)^17, 20^. For the sake of simplicity, we collectively refer to these peripheral neuropathies as Charcot-Marie-Tooth disease (CMTD), which together afflict ~1:2500 individuals, with 70% of cases being due to due to mutations that impact *pmp22* ^17^. The underlying cause of the disease is due to dysmyelination of PNS nerves, which reduces nerve conduction velocity along the peripheral axons. Depending on the causative mutation, CMTD ranges in severity, with symptoms including but not limited to abnormalities of peripheral axons, impaired tendon reflexes, progressive weakness of distal musculature, muscle cramping, and abnormal gait. Patients with a severe phenotype can be confined to a wheelchair, experience chronic pain, and/or be afflicted with blindness and auditory loss ^21–23^. There is presently no treatment for CMTD beyond symptom management^22^.

**Figure 1.**
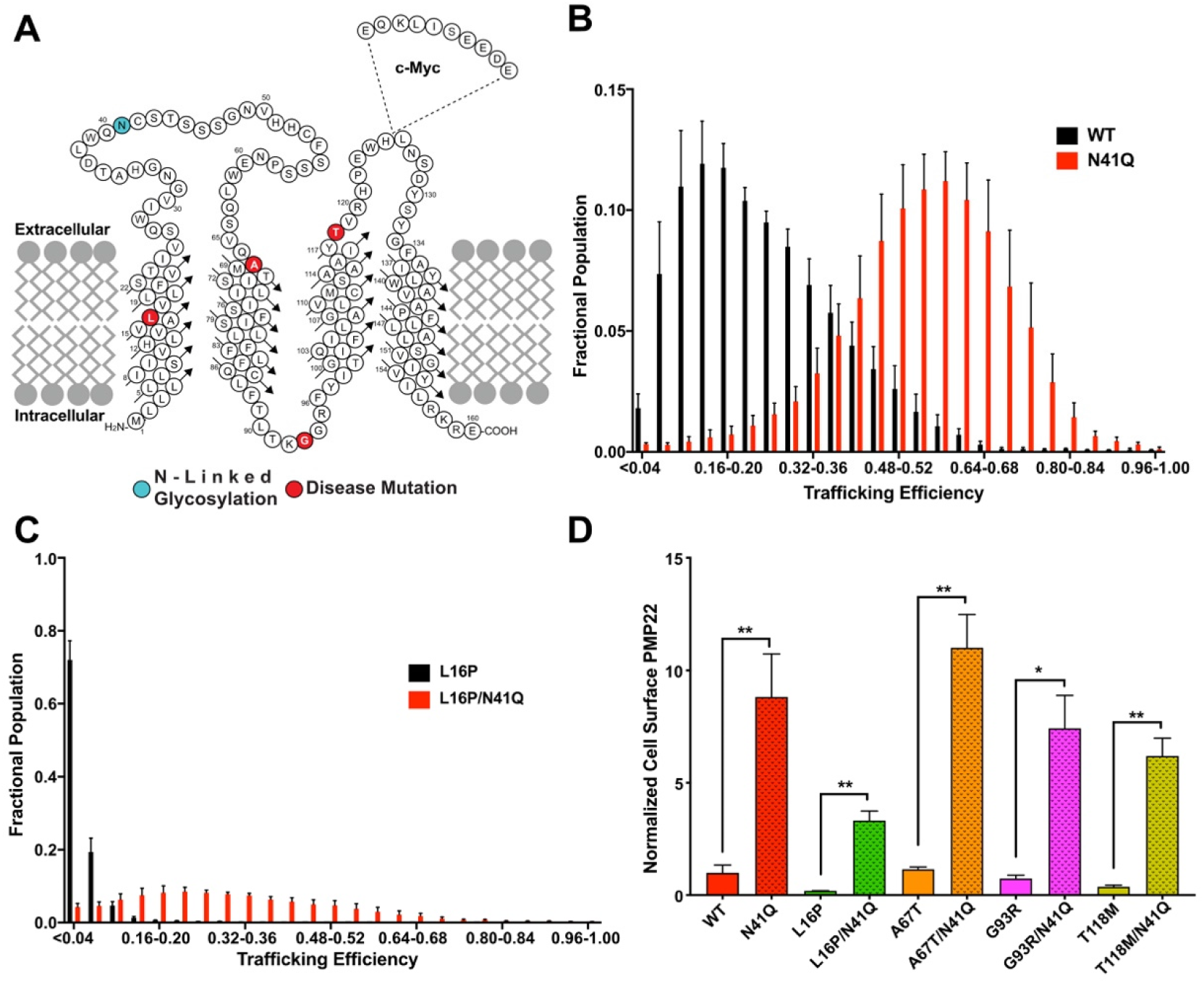
N-Glycosylation limits forward trafficking of PMP22. **(A)** Topology diagram of PMP22 in a membrane. The sequential location of the c-myc tag is shown. Disease variant sites for mutants examined in this study are highlighted in red and the site of N-linked glycosylation is shown in cyan. (**B)** Population distribution of PMP22 trafficking efficiencies measured in individual HEK293 cells for WT (black) and N41Q (red) and for **(C)** L16P (black) and L16P/N41Q (red). Measurements reflect the results from 5 biological replicates with 2500 cells measured per replicate. Error bars represent standard deviations (SD) of the replicates. **(D)** Normalized cell surface expression of PMP22 variants and their glycosylation deficient counterpart. Values reflect outcomes from 5 biological replicates with 2500 cells measured per replicate. All values were normalized to WT PMP22 cell surface expression based on results from paired biological replicates. Error bars represent SD of the replicates. Student’s t-test was used for statistical analysis. *=p<0.05, **=p<0.01.

The most common form of CMTD (type 1A) is caused by overproduction of PMP22, due to a heterozygous duplication of chromosome fragment 17p.11-2.12, resulting in trisomy (three copies) of the *pmp22* gene^20^. One hypothesis for why WT PMP22 overexpression causes disease is that increased production of the protein results in oversaturation of ERQC, leading to accumulation of misfolded protein and resulting toxicity and/or cell stress ^24–25^. This seems especially plausible in light of data indicating that even under healthy conditions, PMP22 is prone to misfold, with only 20% of newly expressed protein trafficking to the cell surface ^24, 26–28^. Mutant forms of PMP22 are known to traffic even less efficiently than WT, consistent with the notion that peripheral neuropathies associated with these PMP22 variant are also the consequence of defects in PMP22 trafficking.

We have previously carried out studies to elucidate the molecular defects in PMP22 that cause it to be mistrafficking-prone. Biophysical studies of PMP22 in detergent micelles revealed that the WT protein is only moderately stable, with the folded conformation favored over unfolded protein by only 1.5 ± 0.1 kcal mol^-127, 29^. While undoubtably more stable in native membranes, marginal stability of PMP22 likely accounts for the known inefficient folding and trafficking even of the WT form to the PM. Disease mutant forms of PMP22 were shown to be even less stable than WT. Indeed, quantitative cell trafficking measurements for this same panel of mutants revealed linear relationships between PMP22 surface trafficking efficiency and patient nerve conduction velocities, between PMP22 stability and surface trafficking efficiencies, and between PMP22 stability and patient nerve conduction velocities^27^. PMP22-linked CTMD appears to be a disease that resembles cystic fibrosis in the sense that the vast majority of disease cases are caused by folding defects and mistrafficking^1, 30^. ERQC is evidently attuned to be able to assess the conformational stability of PMP22.

Exactly how PMP22 folding is monitored and managed in the ER is not well understood. Common ER-resident chaperones involved in the folding of many soluble proteins, such as the heat shock protein (HSP) 70 binding immunoglobin factor (BiP), calreticulin, or the thiol-isomerase ERp57, do not appear to be important for the maturation of PMP22^31–33^. On the other hand, the lectin chaperone calnexin (CNX) has been shown to engage WT and disease variants of PMP22 ^31–35^. Moreover, data indicates that RER1 can retrieve disease variants of PMP22 from the Golgi complex and return it to the ER^35^. Beyond this, little is understood about the components of ER quality control that engage nascent PMP22 to determine the balance between its forward trafficking, retention in the ER, and targeting for degradation. In this paper, we determine that N-linked glycosylation of PMP22 significantly hinders the forward trafficking of both WT and disease variants of the protein. We also explored how PMP22 is N-glycosylated by the oligosaccharyltransferases A and B (OST-A, OST-B). Furthermore, we used quantitative proteomics and related pathway analysis to identify how the assembly of various forms of PMP22 (WT, non-glycosylated, and disease mutant forms) is differentially managed by ERQC. Finally, we also used CRISPR/Cas9 to generate knockout cell lines to illuminate the roles of several key ERQC proteins in mediating the trafficking of different forms of PMP22.

## Results

### N-Glycosylation limits forward trafficking of PMP22

PMP22, like most proteins that are inserted into the ER, is post-translationally modified via the addition of a 14-sugar complex oligosaccharide (N-glycan) ^36–37^. This N-glycan is added to asparagine residues associated with the sequence motif N-X-S/T (where X is any amino acid except proline). PMP22 contains a single N-glycosylation site, located in its extracellular loop 1 (ECL1) at asparagine-41 (N41; **Figure 1A** cyan). Within the lumen of the ER, changing the identity of the N-glycan through addition or removal of monosaccharides can trigger binding of client proteins by a subset of folding-assistive proteins known as lectin chaperones^37–38^. Previous work has shown that for PMP22 the N-glycan may play a modest role in oligomer stability ^39^ but does not seem to affect protein function. We have also recently shown that N-glycosylation of PMP22 does not affect its membrane phase preference for cholesterol-rich ordered phase domains^40^. Here we tested the impact on human PMP22 trafficking of mutating N41 to a glutamine (N41Q), thereby rendering PMP22 glycosylation deficient.

Using single-cell flow cytometry-based to directly quantitate both cell surface and internal (mis-trafficked) levels of PMP22 in individual cells, we measured the trafficking efficiencies (the amount of cell surface PMP22 over total expressed PMP22) for the WT versus N41Q mutant forms PMP22 in HEK293 cells (**Figure 1B**). WT PMP22 displayed a trafficking efficiency of 18.6 ± 5.2% (mean ± standard deviation; **Figure 1B**, black) which corroborates with previous values obtained using the same assay with Madin-Darby Canine Kidney cells and also previous measurements using sciatic nerve lysates^24, 26–27^. Remarkably, we found that N41Q PMP22 trafficked to the cell surface with a nearly 3-fold greater efficiency of 53 ± 7% (**Figure 1B**, red). We then examined a highly destabilized and disease-causing variant of PMP22, L16P, to see if glycosylation also affected its trafficking efficiency (**Figure 1C**). The severe CMTD L16P mutation in PMP22 causes a kink in TM1 that significantly destabilizes the protein and causes the majority of the protein to be retained intracellularly^27, 41–43^. Our experiments reflect these previous observations, as L16P PMP22 was seen to surface-traffic with only 3 ± 1% efficiency (**Figure 1C,** black). However, when the glycosylation site was removed in this variant (L16P/N41Q) the trafficking efficiency increased to 34 ± 10%, a more than 10-fold increase(**Figure 1C,** red). We also observed a pronounced increase in surface trafficking efficiency upon removal of N-linked glycosylation for three additional CMTD PMP22 mutants: one known to have WT-like stability (A67T) and two with stability intermediate between WT and the unstable L16P (G93R and T118M) (**Supplemental Figure 1**)^27^.

In ^39^, the authors explored the role of N-linked glycosylation on WT PMP22 stability, oligomerization and localization. They observed no differences in PMP22 localization between the glycosylated and nonglycosylated form using cell surface biotinylation or confocal microscopy. However, the authors did note that they could have missed changes in PMP22 distribution due to limitations in the techniques then available. The trafficking assay employed in this study is both robust and quantitative, explaining why this change in PMP22 trafficking efficiency owing to N-linked glycosylation was not previously detected.

Deconvolution of the data to examine total, internal, and cell-surface concentrations of PMP22 provides additional insight into the role of N-glycosylation in PMP22 trafficking (**Figure 1D, Supplemental Figure 2**). For WT and all PMP22 disease variants, we noted a drastic and significant increase in cell surface expression for glycosylation-deficient variants compared to normally glycosylated isoforms (**Figure 1D**). Conversely, we noticed smaller changes in the amount of internally trapped PMP22 (**Supplemental Figure 2A**) and total expression levels (**Supplemental Figure 2B**) when we compared glycosylated versus non-glycosylated PMP22 variants. This data led us to hypothesize that N-linked glycosylation plays a central role as a bottleneck to forward-trafficking of PMP22, limiting the amount of protein that reaches the cell surface.

### Mechanism of PMP22 glycosylation

In light of the observation that N-linked glycosylation limits PMP22 forward trafficking, we sought to uncover the pathway by which this modification occurs. In mammalian cells, the oligosaccharyltransferase (OST) complex catalyzes the transfer of a preassembled oligosaccharide from a dolichol pyrophosphate-linked donor onto the target protein^36–37^. Mammalian cells express two OST complexes with different catalytic subunits (STT3A and STT3B), a shared set of non-catalytic subunits, plus some complex-specific subunits^44^. Complexes containing STT3A (OST-A) are associated with the Sec61-translocon and catalyze canonical co-translational glycosylation as proteins are threaded into the ER^45–46^. Complexes containing STT3B (OST-B) are not associated with the translocon and catalyze glycosylation post-translationally^46^. We sought to uncover which OST complex—OST-A or OST-B—was responsible for mediating PMP22 glycosylation.

We quantified PMP22 glycosylation via western blotting in cell lysates from both HEK293 cells and from HEK293 cells in which STT3A or STT3B had been genetically knocked out using CRISPR/Cas9 (**Figure 2**)^44^. WT PMP22 separates into three distinct bands on an SDS-PAGE gel (**Figure 2A**), where the identities of each can be confirmed via comparison with gel patterns following treatment of samples with different glycosidases. The lower band corresponds to non-glycosylated PMP22 (the band is unchanged upon treatment with glycosidases), the middle band corresponds to ER-resident PMP22 (as identified by its disappearance upon treatment with EndoH), and the top smear corresponds to post-ER PMP22 (as identified by its resistance to EndoH and disappearance upon treatment with PNGase). Quantification of the fraction of PMP22 that is glycosylated in these cell lines from three independent biological replicates revealed that 69 ± 2% (mean ± standard deviation) of total WT PMP22 is glycosylated in HEK293 cells, 63 ± 3% is glycosylated in STT3A knock-out (KO) cells, and 48 ± 3% is glycosylated in STT3B KO cells (**Figure 2A**). These results suggest that while both OST complexes are able to glycosylate WT PMP22, post-translational glycosylation mediated by OST-B appears to be the predominant pathway, as the STT3B KO cell line caused a significant loss in PMP22 glycosylation, while the level of glycosylation in the STT3A KO cell line remained similar to normal HEK293 cells. To confirm this, we also quantified WT PMP22 glycosylation in MagT1/Tusc3 double KO cell lines. MagT1 and Tusc3 are accessory proteins found exclusively in the OST-B complex^47^. In this cell line, 40 ± 7% of WT PMP22 was glycosylated, confirming OST-B as the predominate pathway of N-glycosylation for WT PMP22.

**Figure 2.**
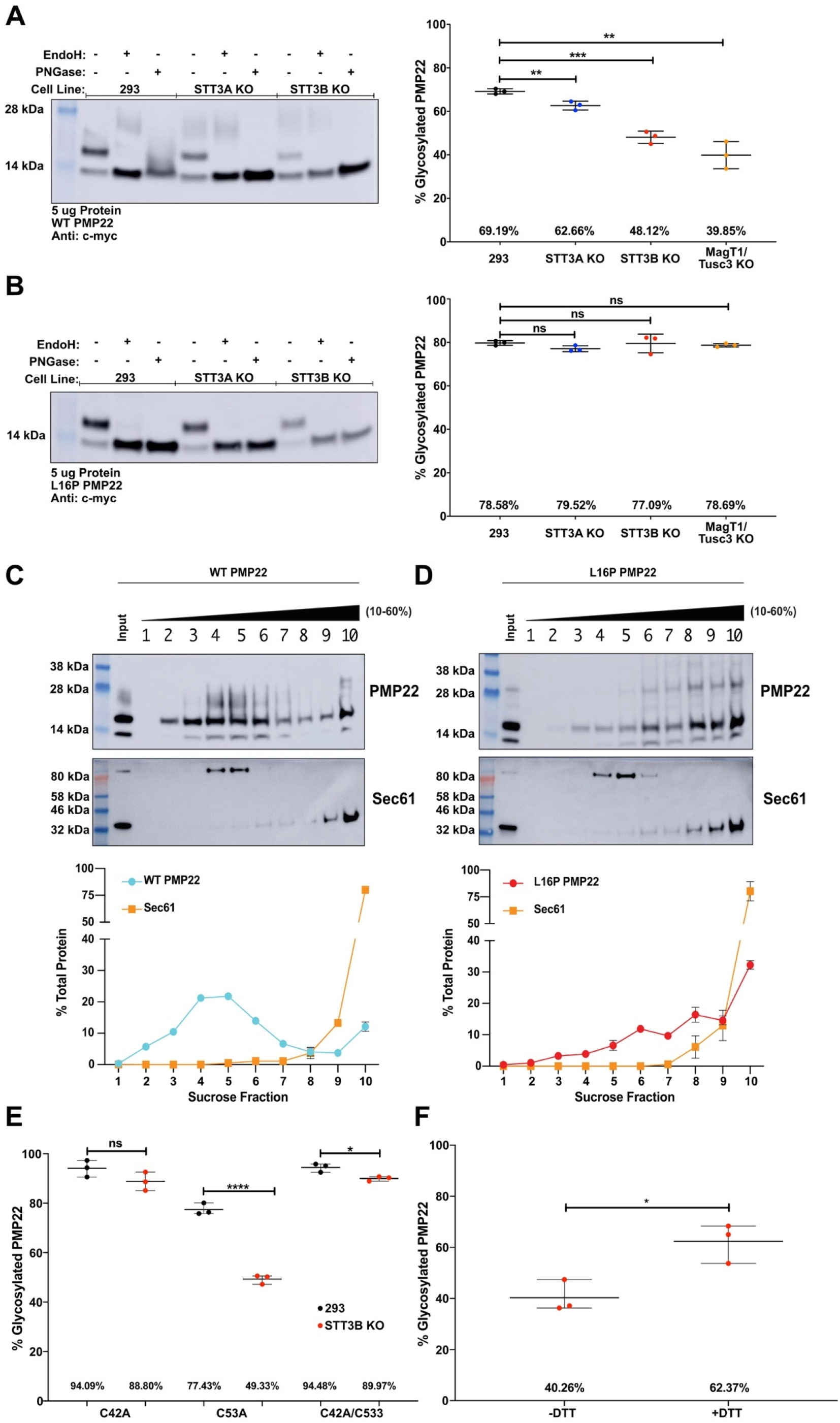
Mechanism of PMP22 glycosylation. **(A, B)** Western blots showing WT PMP22 **(A)** or L16P **(B)** PMP22 glycosylation in HEK293 cells, STT3A KO HEK293 cells, or STT3B HEK293 cells. Glycosidase treatments with EndoH or PNGase was used to confirm the identity of the bands. Quantified levels of glycosylated PMP22 from 3 independent biological replicates are shown on the right. Uncut western blots are shown in **Supplemental Figure 3 (C,D)**. Sucrose density fractionation of cell lysates containing WT **(C)** or L16P (**D)**. Representative blots showing the distribution of PMP22 and Sec61 are shown, with quantified levels of the two proteins from three independent biological replicates shown below. Uncut western blots are shown in **Supplemental Figure 5**. **(C)** Quantified levels of glycosylated C42A, C53A, or C42A/C53A PMP22 in HEK293 (black) or STT3B KO HEK293 (red) cells from three independent biological replicates. **(F**) Quantified levels of glycosylated WT PMP22 in STT3B KO HEK293 cells that were either not treated or treated with 2 mM dithiothreitol (DTT) for 2 hours prior to cell lysis. All error bars represent standard deviations and, if not shown, were too small to be visualized. Students t-test was used for all statistical analyses. ns = not significant, *p<0.05, **p<0.01, ***p<0.001, ****p<0.0001.

We next quantified the levels of L16P PMP22 glycosylation in these four cell lines (**Figure 2B**). In HEK293 cells 80 ± 2% of L16P PMP22 was glycosylated, while 77 ± 2% was glycosylated in STT3A KO cells, 80 ± 5% was glycosylated in STT3B KO cells, and 79 ± 1% of was glycosylated in MagT1/Tusc3 double KO cells. This contrasts with the WT PMP22 results, suggesting that L16P PMP22 has no preference for glycosylation via OST-A vs. OST-B. We also quantified the levels of glycosylation in HEK293, STT3A KO and STT3B KO cell lines for A67T, G93R, and T118M PMP22 (**Supplemental Figure 4**) and found similar results as for L16P PMP22. These results suggest that unlike with the moderately stable WT protein, less stable and misfolding-prone CMTD variants of PMP22 can be glycosylated equally well co- or post-translationally. This suggests that for these variants of PMP22, co-translational glycosylation via OST-A plays the predominant role in PMP22 glycosylation, since the protein will be exposed to OST-A before it reaches OST-B.

Why is glycosylation of WT PMP22, but not CMTD variants, sensitive to the loss of OST-B? One hypothesis is that the conformational instability of the CMTD mutants causes these proteins to remain associated with the translocation machinery and therefore in proximity to OST-A longer than for WT PMP22. As seen in **Figure 2A**, a significant portion of WT PMP22 is still glycosylated even in the absence of STT3B suggesting that WT PMP22 is still a substrate, albeit a suboptimal one, for OST-A glycosylation. If the CMTD variants are ‘stuck’ on the translocon they would be retained in close proximity to OST-A for a longer period of time than the WT protein, allowing for more efficient glycosylation. To test this hypothesis, we monitored direct association of WT and the unstable L16P form of PMP22 with the Sec61 translocon. To this end, we performed sucrose density fractionation of PMP22 containing-cell lysates^48^. Western blotting was used to locate PMP22 as well as the Sec61 subunit of the translocon^4–6, 45^ in the gradient. As seen in **Figure 2C,** WT PMP22 preferentially partitioned in fractions 3-6 with a second minor population partitioning in fraction 10. Sec61 predominantly partitioned in fractions 9 and 10, indicating that the majority of WT PMP22 is not associated with the translocon, as expected for this relatively well-folding form of the protein. **Figure 2D** shows the fractionation of L16P PMP22. Unlike WT PMP22, L16P PMP22 partitions into a higher density fraction with a significant amount co-partitioning with Sec61. This suggests that more L16P PMP22 is retained at the translocon, explaining why its glycosylation is insensitive to STT3B KO. L16P PMP22 appears to be insensitive to STT3B KO due to its increased association with the translocon and the accompanying OST-A complex.

The second question arising from **Figure 2A** is: why does a significant fraction of WT PMP22 escape OST-A glycosylation as it is threaded into the ER lumen? It has been observed that glycosylation sequences with a cysteine residue at the N+1 position relative to the glycosylation site (-N-C-S/T sequences) tend to be skipped by OST-A^47^. Human PMP22 contains a cysteine residue adjacent to the glycosylation site at amino acid residue 42 (C42). To test the importance of C42 in PMP22 glycosylation, we made an alanine mutation (C42A) and measured PMP22 glycosylation in HEK293 cells and STT3B KO cells. We also mutated the other solvent exposed cysteine in PMP22 to an alanine (C53A) and also made the double cysteine to alanine mutations (C42A/C53A). **Figure 2E** shows that like WT, the glycosylation of C53A PMP22 is sensitive to the loss of the OST-B complex, leading to a reduction in glycosylation (77 ± 3% glycosylated in HEK293 cells and 49 ± 2% glycosylated in STT3B KO cells). This suggests that C53A PMP22 is still predominately glycosylated post-translationally via OST-B.

However, in the single C42A or double C42A/C53A PMP22 mutants, glycosylation was no longer sensitive to STT3B KO. C42A PMP22 was glycosylated 94 ± 4% in HEK293 cells and 89 ± 4% in STT3B KO cells while C42A/C53A PMP22 was glycosylated 95 ± 2% in HEK293 cells and 90 ± 1 % in STT3B KO cells. This data suggests that these mutants can now be glycosylated via OST-A. Interestingly, C42A and C42A/C53A PMP22 exhibited higher levels of glycosylation in HEK293 cells than WT PMP22, implying that the C42 helps make WT PMP22 a suboptimal substrate for STT3A glycosylation.

Structural studies comparing OST-A and OST-B complexes do not show significant differences in the active sites that could explain the differences in substrate specificity^49^. While it is unknown if C42 and C53 are involved in an intramolecular disulfide bond in PMP22^18^, the fact that the C53A mutation did not affect PMP22 glycosylation ruled out the possibility that a disulfide bond between C42 and C53 impedes PMP22 glycosylation by STT3A. However, we hypothesized that the redox state of C42 may be responsible for making PMP22 a suboptimal substrate for OST-A. To test this hypothesis, we measured PMP22 glycosylation in STT3B KO cells that had been treated with small amounts (2 mM) of the reducing agent dithiothreitol (DTT) for 2 hours prior to cell lysis (**Figure 2F**). This short treatment is not long enough to induce changes in the ER proteome due to activation of the unfolded protein response^50^. We observed an increase in PMP22 glycosylation in STT3B KO cells from 40 ± 7% in untreated cells to 62 ± 8% in cells treated with DTT. While this short treatment did not restore the levels of WT PMP22 glycosylation to that observed in HEK293 cells (**Figure 2A**) it did cause a significant increase. This result combined with that observed in **Figure 2E** suggests that the efficiency of N-glycosylation of WT PMP22 by OST-A is dependent on the thiol redox potential of the lumen of the ER in a way that is dependent on PMP22 C42, with a more reducing environment resulting in higher efficiency of N-glycosylation by OST-A. Because C42 does not form a disulfide bond with C53, this suggests the possible involvement of some other ER-resident thiol redox protein, perhaps via transient disulfide bond formation with C42 of PMP22.

### PMP22 trafficking in response to loss of glycosylation machinery

We next tested what happens to PMP22 trafficking when specific mechanisms of glycosylation were inhibited. We measured PMP22 trafficking in STT3A and STT3B KO cell lines (**Figure 3,** horizontal and vertical stripped graphs respectively). Additionally, we measured PMP22 trafficking under conditions in which both complexes or STT3B only were inhibited for 24 hours prior to the experiment (**Figure 3** checkered and dotted plots respectively). NGI-1, a partial inhibitor of both STT3A and STT3B^51^, resulted in decreased WT PMP22 glycosylation to 12 ± 1 % (**Supplemental Figure 8**). C19, a specific inhibitor of STT3B^52^, reduced WT PMP22 glycosylation to 48 ± 2%, similar to what was observed in the STT3B KO cells (**Supplemental Figure 8**).

**Figure 3.**
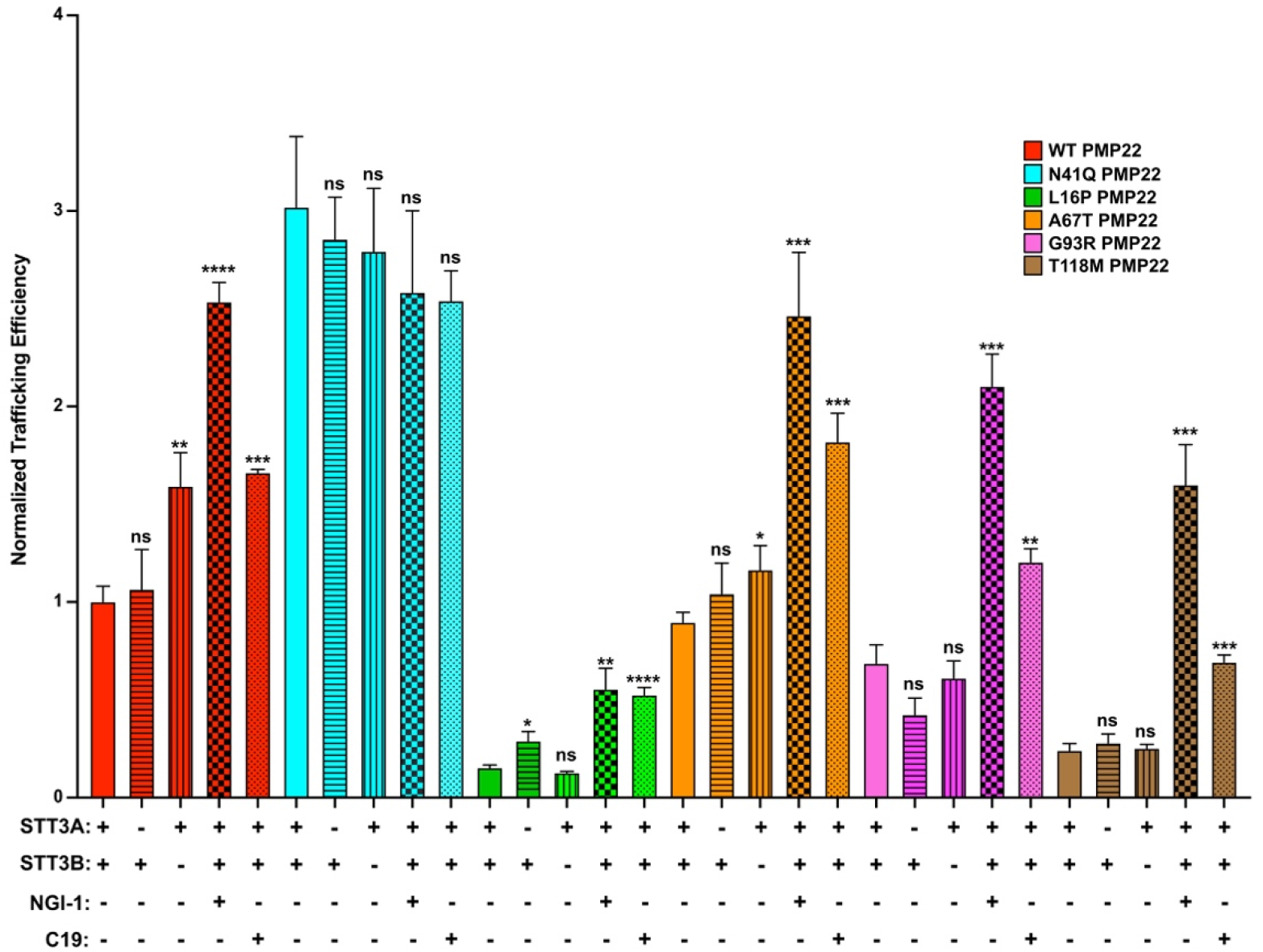
PMP22 trafficking efficiencies measured in the presence of glycosylation inhibition. Normalized trafficking efficiencies of PMP22 variants in HEK293 cells (solid color), STT3A KO HEK293 cells (horizontal stripes), STT3B KO HEK293 cells (vertical stripes), HEK293 cells treated for 24 hours with 10 μM NGI-1 (checkered, OSTA/OSTB inhibitor) or HEK293 cells treated for 24 hours with 10 μM C19 (dots, OSTB inhibitor) from three independent biological replicates. All efficiency values were normalized to WT PMP22 trafficking in untreated HEK293 cells in paired biological replicates. Error bars represent SD. Student’s t-test was used to compare the trafficking efficiencies in untreated cells to that in KO cell lines or in cell lines treated with the inhibitors. ns = not significant, *p<0.05, **p<0.01, ***p<0.001, ****p<0.0001.

In **Figure 3** we measured PMP22 trafficking efficiencies under these various conditions. All efficiency values were normalized relative to WT PMP22 in paired experiments using either untreated HEK293 cells (experiments with STT3A or STT3B KO cells) or HEK293 cells treated with DMSO (experiments with NGI-1 or C19). As expected, STT3A KO had no effect on WT PMP22 trafficking efficiency, confirming its minor role in mediating WT PMP22 glycosylation. However, in STT3B KO cells WT PMP22 exhibits a 1.6 ± 0.4-fold increase (mean ± standard deviation) in trafficking efficiency. Treatment of HEK293 cells with NGI-1 further increased WT PMP22 trafficking efficiency 2.5 ± 0.2-fold over untreated cells, similar to the trafficking efficiency of N41Q PMP22. Treating cells with the STT3B specific inhibitor C19 caused a 1.7 ± 0.1-fold increase over untreated cells. WT PMP22 exhibited a significant increase in trafficking efficiency in cells treated with C19 or in STT3B KO cells, but not in STT3A KO cells. These results confirm that OST-B is the major facilitator of WT PMP22 glycosylation.

Since N41Q PMP22 is not glycosylated we expected no changes in it trafficking efficiency under any of these OST inhibitory conditions. As predicted, none of these experimental conditions significantly altered the trafficking efficiency of N41Q PMP22 (**Figure 3**). This result serves as an internal control for our experiments by showing that neither use of OST KO cell lines nor pharmacological inhibition of OST caused glycosylation-independent global changes for this mutant in its protein folding quality control and surface trafficking efficiency.

Inhibition of both OSTs via NGI-1 caused increases in trafficking efficiency for all CMTD PMP22 variants studied (**Figure 3**). However, neither STT3A nor STT3B KO cell lines dramatically altered the trafficking efficiencies of these variants. This makes sense in light of the data in **Figure 2B** and **Supplemental Figure 4** showing that KO of either STT3A or STT3B did not reduce the glycosylation levels of these PMP22 variants to the same extent as for WT PMP22. Interestingly, specific inhibition of STT3B with C19 caused an increase in PMP22 trafficking efficiencies of all three CMTD mutants. Our data clearly indicates a significant fraction of PMP22 is glycosylated post-translationally by STT3B and that inhibition of this process, either genetically or pharmacologically, increases PMP22 trafficking efficiency.

### Identification of PMP22 interactor proteins

We next sought to uncover proteins responsible for limiting or promoting PMP22 trafficking. Prior to this work, the only ERQC proteins that have been experimentally shown to interact with PMP22 are CNX and RER1^31–33, 35^. In order to discover novel PMP22 interactors, we turned to co-immunoprecipitation (co-IP) and liquid chromatography-mass spectrometry/mass spectrometry (LC-MS/MS) based proteomics. We expressed myc-tagged WT and mutant forms of PMP22 in HEK293 cells, immunopurified the protein using magnetic beads conjugated to myc antibodies under gentle lysis conditions, and used shotgun proteomics to identify proteins that immunopurified with PMP22. Tandem mass tag (TMT) labeling was used to quantitatively compare interactions across multiple samples^53–54^. We compared interactions with WT PMP22 versus N41Q PMP22 to uncover glycosylation specific interactions. Furthermore, WT versus L16P PMP22 measurements were used to uncover interactions that depend on protein stability. Proteins co-purified with tagged PMP22 were also compared to proteins that were co-purified from a mock lysate (cells expressing untagged PMP22) to remove non-specific background. **Figure 4** displays examples of the PMP22 interacting proteins identified by our screen. The full results from this screen are found in **Supplemental Table 1**.

**Figure 4.**
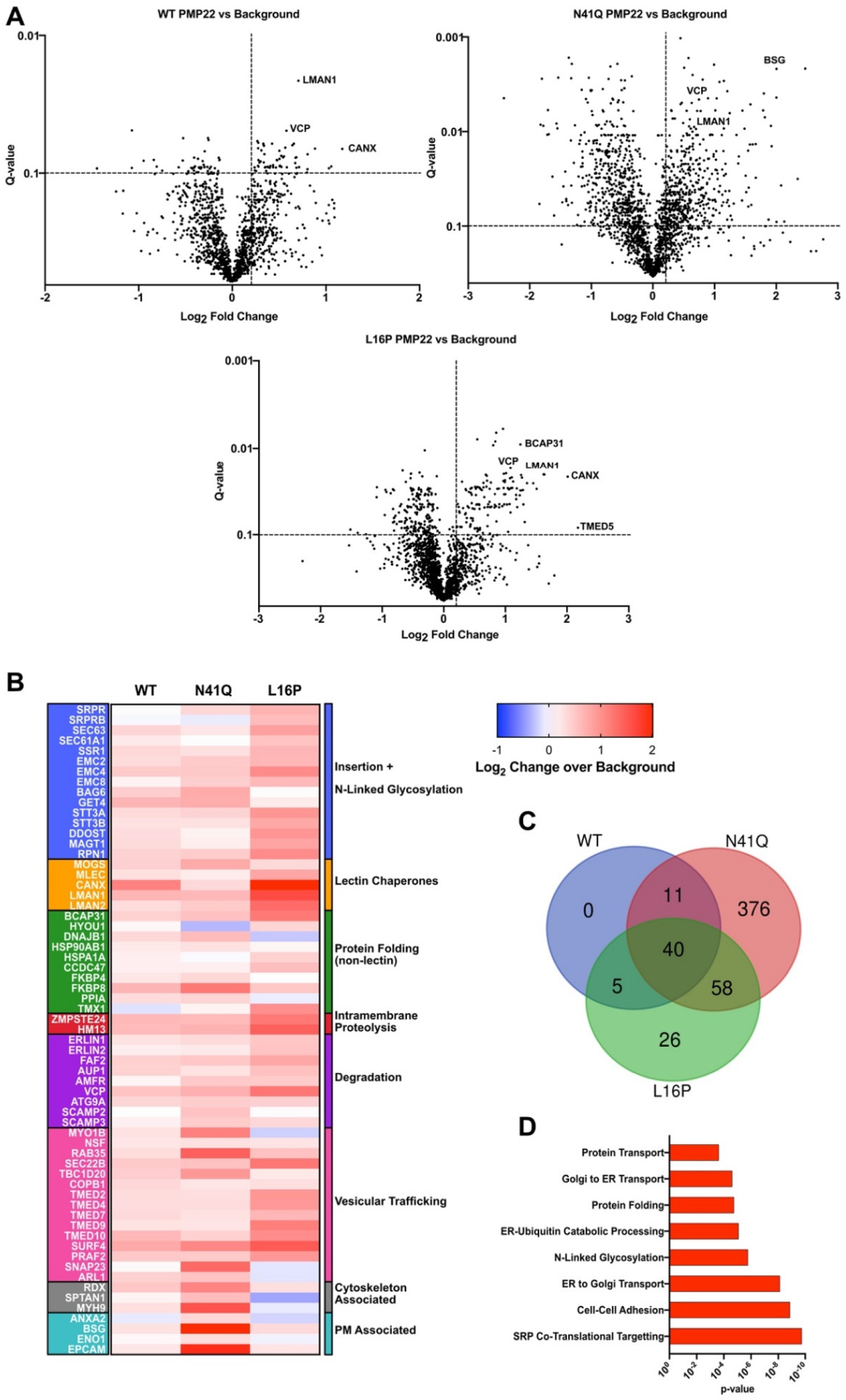
Proteomics to uncover novel PMP22 interacting partners. Co-immunoprecipitation and LC-MS/MS proteomics was used to identify protein that interact with PMP22. **(A)** Volcano plots for WT, N41Q, and L16P PMP22 showing interacting proteins that were identified from 5 biological replicates. Log2 fold change of the identified protein in the WT PMP22 sample as compared to samples obtained from IPs with untagged PMP22 are shown on the x-axis and the statistical significance for the interactions (Q-value) from the multiple replicates is shown on the y-axis. Selected interactions are identified in which dotted lines separate the data into quadrants with the upper righthand quadrant signifying PMP22 interactors. **(B)** Heat-map showing quantified interactions for WT, N41Q, or L16P PMP22 over background. The heat map is arranged temporally in regard to PMP22 biogenesis. The top of the map identifies interactions predicted to occur early in biogenesis and the bottom of the map shows interactions predicted to occur later in biogenesis. All interactions had a Log2-fold change over background >0.2, Q-value <0.1 and were identified in multiple biological replicates for at least one PMP22 variant. **(C)** Venn-diagram showing overlap of interacting partners identified for WT, N41Q, or L16P with a Log2-fold change over background >0.2, Q-value <0.1. **(D)** Pathways enrichment analysis showing the GO_Biological Processes enrichment from the identified PMP22 interactors. A Fisher Exact p-value test was used to calculate the p-values for the GO_term enrichment.

Overall, we identified 56 unique proteins that appear to interact either directly or indirectly with WT PMP22 (**Figure 4A**). Additionally, 485 and 129 proteins appear to interact with N41Q and L16P PMP22 respectively (**Figure 4A, C**). We used the DAVID bioinformatic database to initially group these interactions based on gene ontology (GO) terms followed by a manual curation of the interactions based on literature analysis, and composed the diagram shown in **Figure 4B** to visualize interactions known to be involved either in protein biogenesis, quality control, or other possible functions ^55–56^. Importantly, the TMT quantification allows for direct comparisons of partner protein abundances across the different PMP22 variants. The table is organized temporally with respect to the start of biogenesis at the translocon, with interactions predicted to occur earlier in PMP22 biogenesis shown at the top (insertion and N-linked glycosylation) and those expected to occur later (plasma membrane localized) shown at the bottom. It was reassuring that we were able to identify CNX in our screens and that the ordered affinities of CNX for different forms of PMP22 were L16P>WT>N41Q—as supported by previous literature results ^31–34^. **Figure 4C** shows a Venn diagram of the identified interactions for all three PMP22 variants. In all cases putative interactors were observed at levels with a log2-fold change of >0.2 over mock conditions. Moreover, they were identified in multiple biological replicates and exhibited a Storey Q-value <0.1 compared to mock samples (**Supplemental Table 1**). **Figure 4D** shows biological processes that were the most highly represented based on the results of pathway enrichment analysis using the identified interactors.

We noticed a couple of key trends when the data was organized in the interaction network implied by **Figure 4B**. First, N41Q PMP22 had significantly higher interactions with proteins later in the pathway (cytoskeleton associated, and plasma membrane associated) than WT or L16P PMP22. This observation makes sense in light of the results presented in **Figure 1**. Since N41Q PMP22 traffics to the cell surface much more efficiently than WT PMP22 it is expected that its interactions with proteins at later stages in the trafficking pathway should be more abundant. A second observation from the proteomic analysis is that the folding defect caused by the L16P mutation in PMP22 seems to be recognized very early in biogenesis. L16P PMP22 shows much higher interactions than WT or N41Q PMP22 with proteins associated with the ‘insertion and N-linked glycosylation’ and ‘lectin chaperone’ stages. This suggests that more and/or longer-lived binding events occur early on for this variant, confirming the results of the OST experiments presented earlier in the Results. These results support our hypothesis that unstable variants of PMP22 such as L16P tend to get retained on the translocon compared to WT PMP22, as L16P PMP22 interacted more strongly with Sec61 than WT or N41Q PMP22 (**Figure 4B**).

### N-Glycan recognizing chaperones involved in PMP22 trafficking

Some of the putative interactors uncovered in our proteomic screen are known to bind client proteins in an N-glycan dependent manner (lectin chaperones). We set out to validate the role of these interactors in mediating PMP22 trafficking. We also were interested in the non-lectin chaperone RER1 in light of a previous report that it is involved in ERQC for PMP22^35^. To validate whether these proteins do indeed function in PMP22 trafficking, we generated CRISPR/Cas9 clonal KO cell lines (**Supplemental Figure 9**) and quantitated PMP22 trafficking efficiency therein. We focused on four potential mediators of PMP22 trafficking: CNX, lectin mannose-binding protein 1 (LMAN1, also known as ERGIC53), UDP-glucose:glycoprotein glucosyltransferase 1 (UGGT1), and RER1. CNX and RER1 are the only previously-identified chaperones that engage PMP22^31–33, 35^. CNX is believed to retain PMP22 in the ER to promote folding, while RER1 is a sorting receptor in the Golgi and has previously been shown to function in retrograde trafficking of two variants of PMP22: L16P and G150D. LMAN1 was one of the strongest interactors to come out of our screen (**Fig. 4A-B & Supplemental Table 1**) and is a mannose specific lectin that has previously been shown to promote maturation of glycoproteins from the ER to the Golgi complex^57–59^. We hypothesized that LMAN1 might be responsible for promoting maturation of PMP22 in a glycan specific manner. UGGT1 is a main component of the CNX cycle and has been proposed to act as its main folding sensor. We did not identify UGGT1 in our screen, suggesting short lifetimes for UGGT1-PMP22 complexes. However, since PMP22 was seen to interact strongly with CNX and since UGGT1 is a main component of the CNX cycle we hypothesized a likely interaction. UGGT1 is thought to act either by re-glucosylating folding-immature client proteins to cause reengagement with CNX or by recognizing polypeptides as folded, with a “no glucosylate” decision then being made that allows the client protein to forward-traffic beyond the ER^10, 60^. We hypothesized that a loss of UGGT1 would cause an increase in PMP22 maturation in a glycosylation dependent manner.

**Figure 5** shows PMP22 trafficking efficiencies of WT, N41Q and L16P PMP22 in the four KO cell lines compared to HEK293 cells. The data is presented as violin plots, which show the population distribution of PMP22 trafficking efficiencies in individual cells from three independent biological replicates measuring >2000 cells per replicate. The horizontal white lines separate the data into quartiles. (Data for A67T, G93R, and T118M PMP22 can be found in **Supplemental Figure 10**).

**Figure 5.**
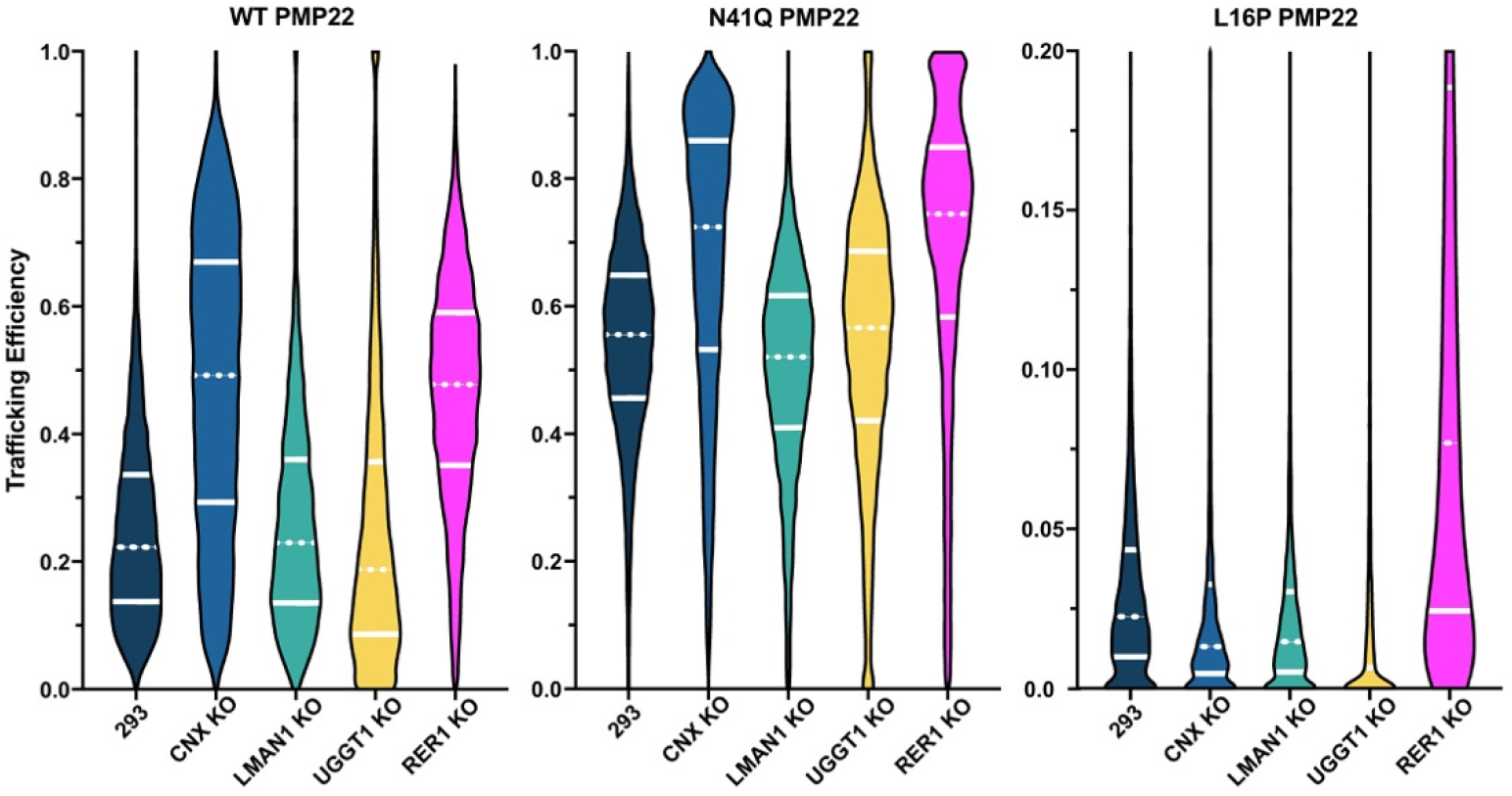
PMP22 trafficking efficiencies in CRISPR/Cas9 KO cells of potential ERQC interactors. CRISPR/Cas9 was used to generated KO cells of selected potential proteins involved in mediating trafficking effects for PMP22 (**Supplemental Figure 10)**. Violin plots showing population distributions of WT, N41Q and L16P PMP22 trafficking efficiencies from three biological replicates are shown. Data was collected in HEK293 cells (navy), CNX KO HEK293 cells (blue), LMAN1 KO HEK293 cells (green), UGGT1 KO HEK293 cells (yellow) and RER1 HEK293 KO cells (pink). White lines in the population distributions separate the data into quartiles, the dashed line in each case represents the median. Pairwise Kolmogorov–Smirnov test was used to compare population distributions.

KO of CNX caused a dramatic, and significant, increase in PMP22 surface trafficking efficiency for both WT and the glycosylation-deficient N41Q PMP22 **Figure 5**. Relative to HEK293, the increase for WT is from a median trafficking efficiency of 22% to 49%. For N41Q the increase in median trafficking efficiency was from 56% to 72%. The fact that this increase was independent of glycosylation makes sense in light of previous experiments showing that even though CNX is a “lectin chaperone” it can engage PMP22 independent of its glycosylation state^34^. These results are at variance with previous studies in which confocal microscopy was used to show that CNX knockdown did not change the distribution of GFP-tagged PMP22 in cells^35^. This discrepancy may be explained by previous studies using GFP-tagged PMP22 showing that the tag promotes intracellular retention of PMP22 because of the ability of GFP to oligomerize, resulting in PMP22 aggregation^61^. Our results suggest that the modest (ca. 20%) trafficking efficiency by which WT PMP22 surface-traffics is mainly due to intracellular retention by CNX, an activity that CNX evidently carries out using both glycosylation dependent and independent mechanisms.

In contrast to the results for WT, L16P PMP22, showed no significant change in PMP22 trafficking efficiency when CNX was knocked out. This strongly implies that CNX is not the primary folding sensor responsible for retaining the most unstable mutant forms of CNX in the ER. G93R and T118M, which have stabilities intermediate between WT and L16P ^27^ both exhibit modest trafficking enhancement in CNX KO cells that are intermediate between WT and L16P (**Supplemental Figure 10**). A67T, which has WT-like stability, exhibits a dramatic enhancement of surface trafficking that resembles that of WT (**Supplemental Figure 10**). These results again point to CNX as the main determinant of PMP22 surface trafficking efficiency for WT and stable mutants, but suggest that another component of ER quality control may play the role of key gatekeeper for unstable disease mutant forms.

To our surprise, KO of LMAN1 showed no effect on the trafficking efficiency of any of the PMP22 variants under study (**Figure 5, Supplemental Figure 10**). This result has more than one possible interpretation. First, it could be be that the complex formed between PMP22 and LMAN1 does not play a major role in directing the surface-trafficking of PMP22. Alternatively, it could mean that other ER sorting receptors, such as LMAN2, which was also identified in the proteomic screen, are functionally redundant with LMAN1, stepping in as a backup to fulfill the function of this lectin chaperone in PMP22 trafficking when LMAN1 is knocked out.

UGGT1 KO cells showed a very significant decrease in PMP22 trafficking efficiency for WT, L16P, G93R and T118M but not for glycosylation-deficient N41Q PMP22 (**Figure 5, Supplemental Figure 10**). A67T PMP22 exhibited the same median trafficking efficiency in UGGT1 KO cells with a slight downward shift in the population distribution (**Supplemental Figure 10**). It is not surprising that N41Q PMP22 showed no changes in trafficking efficiency in UGGT1 KO cells since this PMP22 variant does not contain an N-glycan moiety for UGGT1 to modify. A possible explanation for these results is that by re-glucosylating the PMP22 N-glycan, UGGT1 both promotes correct folding and also protects PMP22 from premature degradation. This hypothesis is supported by observing a decrease in total cellular WT PMP22 levels in UGGT1 KO cells relative to HEK293 (**Supplemental Figure 11**), suggestive of increased PMP22 degradation in UGGT1 KO cells.

RER1 KO cells showed increases in trafficking efficiencies for WT and all PMP22 variants under study (**Figure 5**). This confirms previous results indicating a role for RER1 in ERQC of PMP22 ^35^. The present results suggest that RER1 functions as a glycosylation-independent Golgi-to-ER retrograde transporter for both WT and disease mutant forms of PMP22, a function that reflects a role for RER1 in preventing the escape of folding-immature proteins from the ER. This result expands the role of RER1 as functioning not only in retrograde transport of disease variants of PMP22 but also in retrograde transport of the WT protein.

Overall, these studies show that LMAN1 does not have an obvious role in PMP22 ERQC. CNX plays a critical and at least partly glycosylation-independent role in retaining WT and relatively stable mutant forms of PMP22, but plays a less decisive role for unstable mutant forms of the protein even though it does avidly engage these proteins. RER1 plays a glycosylation-independent role in retaining WT and all mutant forms regardless of stability. Finally, UGGT1 plays a glycosylation-dependent role in retaining WT and PMP22 mutants in the ER and also protects them from degradation.

## Discussion

### N-Glycosylation limits forward trafficking of human PMP22

Under healthy conditions, WT PMP22 traffics to the PM with a modest efficiency of ~ 20%^24, 26–28^. Disease mutant forms of the protein usually traffic with even lower efficiency, in some cases approaching 0%^27, 42^. A key result of this work is that elimination of the single N-glycosylation site in PMP22 (N41) results in several-fold increases in the surface trafficking efficiencies of both WT and disease mutant forms of the protein (**Figure 1**). While N-glycosylation is often associated with promoting molecular recognition, protein stability, or protein solubility^10, 36–38^, it clearly plays a limiting role on PMP22 trafficking from the ER to the PM. This reflects the role of protein N-glycan moieties in ERQC, where they serve as a “quality control barcode” to which monosaccharides are added or removed to signal to the ERQC machinery the folding status of the nascent protein as the basis for determining whether that protein will be retained in the ER for further attempts at folding or whether it should be targeted for degradation^36–37, 62–63^. Mutations of N41 have yet to be discovered among the documented CMTD mutant forms of PMP22. Previous studies showed that N-glycosylation is not important for the ability of PMP22 to partition into cholesterol-rich membrane domains ^40^, form specific *trans* interactions with other myelin proteins ^64^, or modulate lipid ultrastructure *in vitro* ^19^. These reports suggest that the role of N-glycosylation in PMP22 is in ERQC of its folding and trafficking.

Even unstable and very poorly trafficking disease mutant forms of PMP22 exhibit a dramatic enhancement in surface trafficking efficiency upon elimination or inhibition of N-glycosylation(**Figure 1C, Supplemental Figure 1**). It is therefore interesting to wonder if inhibition of glycosylation for these CMTD mutant forms of PMP22 could lead to partial restoration of Schwann cell function that might alleviate some CMTD disease progression and symptoms. This remains to be tested. Inhibition of glycosylation to increase trafficking efficiency could also be investigated as a possible therapeutic approach for the HNPP form of CMTD (~20% of all CMTD cases^17, 20^), which is caused by loss of one WT PMP22 allele (haploinsufficient WT/null conditions). Mutations in STT3A and STT3B, the catalytic subunits to the OST-A and OST-B, respectively, are linked to congenital disorders of glycosylation ^65^. Partial inhibition of these glycosylation pathways has emerged as a potential therapeutic approach for treating certain types of cancers^51–52^. These treatments may be worthy of investigation for treatment of certain forms of CMTD.

### Different pathways of glycosylation for WT and CMTD variants of PMP22

N-linked glycosylation is mediated by the OST complexes in one or both of two ways: co-translationally via OST-A or post-translationally via OST-B^44, 46^. Our results show that for WT PMP22, a significant fraction of the nascent protein is primarily glycosylated by OST-B. In contrast, the severely folding defective L16P disease mutant form of PMP22 can be glycosylated to a much higher extent by OST-A (**Figure 2A** and **2B**). This result, coupled with sucrose density fractionation experiments (**Figure 2C** and **2D**), suggests that L16P PMP22 completes translation and then diffuses beyond the translocon complex only slowly compared to WT. The prolonged residency of L16P PMP22 at the translocon would allow the OST-A glycosylation reaction to efficiently glycosylate this mutant before it reaches OST-B. The L16P mutation is located in the 1^st^ transmembrane helix of the protein, where it induces a folding-destabilizing kink in this helix^43^. The structural and folding defects induced by this mutation are evidently manifested even while the downstream protein sequence is still being translated, providing an example of misfolding that occurs at the stage of membrane integration^66–67^.

**Figures 2E** and **2F** show that the redox potential of the ER lumen impacts the cysteine residue adjacent to the glycosylation site (C42), causing PMP22 to be a suboptimal substrate for STT3A. Because mutating cysteine residue 53 to alanine did not cause PMP22 to become a better substrate for STT3A (**Figure 2E**) we can rule out possible disulfide bond formation between cysteine 42 and cysteine 53 as the basis for this phenomenon. Sequence alignment of human PMP22 with homologs from multiple organisms (**Supplemental Figure 13**) shows that the cysteine at N+1 (relative to the glycosylation site) is highly conserved suggesting its importance in PMP22. The cryo-EM structures for the OST-A and OST-B complex showed that the active sites for both catalytic subunits are very similar, ruling out the possibility that C42 sterically hinders glycosylation by STT3A^49^. Our results suggest that that C42 may engage in an intermolecular disulfide bond with an ER luminal protein as it translocated into the ER, leading to cessation of the OST-A reaction at the associated N-glycosylation site. This would explain why pretreatment of cells with DTT to depress the thiol oxidation potential lowered PMP22 sensitivity to STT3B KO. Indeed, several protein disulfide isomerases capable of such an interaction were identified in our proteomic screen (**Supplemental Table 1**). In contrast to OST-A, OST-B complexes contains MagT1 and Tusc3 subunits, which have oxidoreductase capabilities that could reduce this putative intramolecular disulfide bond to enable OST-B mediated N-linked glycosylation^47^. Additional experiments will be required to fully illuminate the role of C42 in the folding and N-glycosylation of PMP22.

### Proteomic analysis reveals that key ERQC decisions for WT PMP22 occur later in the translocon-to-plasma membrane trafficking pathway than for the L16P disease mutant form

Proteomics followed by pathway enrichment analysis was used to examine the interactome of three PMP22 variants: WT, N41Q and L16P to uncover interactions that were glycosylation dependent (WT vs N41Q) or that depend on the conformational stability of the protein (WT vs L16P). For the glycosylation deficient N41Q we found more interactions with proteins late in the biogenesis pathway and localized at the PM as compared to WT or L16P PMP22. Most interestingly, interactors with the severe disease mutant L16P PMP22 were enriched in proteins residing early in the biogenesis pathway at the level of translocon-mediated membrane integration and N-linked glycosylation. This data suggests that the folding defect in L16P PMP22 is detected very early on in the folding pathway, maybe even at the level of membrane integration. Further emphasizing this point, we found that L16P PMP22 co-partitioned in sucrose density fractionation experiments more strongly with the Sec61 component of the translocon than WT PMP22 (**Figure 2C** and **2D**). Additionally, we found that L16P PMP22 had stronger interactions with the lectin chaperones compared to non-lectin chaperones (**Figure 4B)**. For example, L16P PMP22 had a stronger interaction with the lectin chaperone CNX than WT PMP22, confirming an important prior result^31^. This is consistent with the order-of-magnitude increase in forward trafficking when the L16P glycosylation site is mutated (**Figure 1C**). A possible interpretation of this data is that PMP22 is engaged by lectin chaperones prior to reaching key non-lectin chaperones in its biogenesis pathway, which is consistent with a prior study showing that proteins containing N-glycosylation sites within 50 residues of the N-terminus interact with lectin chaperones prior to non-lectin chaperones ^68^.

### Clarification of the roles of CNX, UGGT1, and RER1 in ERQC management of PMP22 folding and trafficking

Four proteins highlighted by the proteomics data were chosen for further exploration of their roles in PMP22 trafficking: two that had previously been shown to interact with PMP22 (CNX and RER1) and two novel potential interactors (LMAN1 and UGGT1). The roles of these proteins in PMP22 folding and trafficking were examined in cell lines in which the alleles encoding these proteins were genetically knocked-out using CRISPR/Cas9.

CNX has previously been reported to monitor the folding status of PMP22 in a manner that can be either or both glycosylation dependent and independent^31–35^. Fontanini et al. showed that CNX can bind both WT and L16P in their glycosylated forms, with the L16P complex being much longer lived than WT^34^. That same study showed that N41Q PMP22 does not bind CNX, whereas the double L16P/N41Q mutant does. Here, it was observed that the trafficking efficiencies of both WT and N41Q PMP22 increased in the CNX KO cell line. However, the mean trafficking efficiency increased by more than 2-fold for WT PMP22 and only about 25% for N41Q PMP22 in CNX KO cells compared to HEK293 cells (**Figure 5**). This data suggests that CNX can interact with PMP22 in both glycan dependent and independent manners with the former eliciting a stronger effect. It is clear that for WT PMP22 calnexin plays a central role in limiting its forward trafficking efficiency.

L16P PMP22 showed no change in trafficking efficiency in the CNX KO cells. To a degree, this result had been previously suggested, as transient knockdown of CNX did not change L16P PMP22 cell surface levels ^35^. On the other hand, this result is surprising in light of our proteomic data (**Figure 4**) and previous observations ^31, 34^ that CNX forms longer-lived interactions with L16P PMP22 than WT PMP22. The CNX KO results do not point to CNX being the major player in causing L16P PMP22 to have a 10-fold lower trafficking efficiency than L16P/N41Q PMP22 (**Figure 1C**). The fact that L16P/N41Q traffics to the cell surface more efficiently than L16P suggests that the decision to retain L16P early in the secretory pathway likely occurs *before* engagement with CNX. Further experiments will be needed to test this hypothesis. The fact that CNX KO did not have a significant effect on L16P PMP22 trafficking could be explained if an unidentified protein is functionally redundant with CNX, normally serving only as a back-player in support of PMP22 folding, but coming to the fore when CNX is removed. One potential candidate for this function is calreticulin, however we did not observe this in our proteomic data and previous reports have shown that it is not involved in PMP22 biogenesis, at least in the presence of calnexin ^31, 33^. There are previous reports of disruption of individual ER chaperones that exhibited only modest consequences due to their considerable redundancy^69^.

UGGT1 is believed to monitor the folding status of glycoproteins upon entry into the ER and also after they have been released from CNX^10–11, 37^. Following engagement with a folding substrate glycoprotein, UGGT1 either catalyzes the addition of a glucose residue onto the N-glycan in order to drive reengagement with CNX or declines to catalyze glucosylation to allow a correctly folded protein to forward traffic beyond the ER^60^. KO of UGGT1 in HEK293 cells caused a decrease in the trafficking efficiency for both WT and most disease variant forms but did not change the trafficking efficiency of N41Q PMP22 (**Figure 5**). Observation that N41Q PMP22 trafficking efficiency was unchanged is as expected since this protein is not glycosylated. In **Supplemental Figure 11,** it is was also shown that there is a decrease in total WT PMP22 levels in UGGT1 KO cells compared to 293 cells. Our hypothesis to explain the data for the WT and disease mutant forms of PMP22 is that engagement of the protein by UGGT1 not only promotes folding of PMP22 in the ER but also helps protects PMP22 from premature degradation via ERAD.

Finally, we explored the effect of RER1 KO on PMP22 trafficking efficiency. RER1 had previously been characterized to play a role in retrieving misfolded L16P and G150D PMP22 disease mutants from the Golgi and returning these proteins to the ER^35^. It thereby appears to play a role in recognizing any misfolded PMP22 that escapes the quality control network. Not surprisingly, RER1 KO caused an increase in the surface trafficking efficiency of L16P PMP22. However, found that both WT and N41Q PMP22 exhibited increased trafficking efficiency in the RER1 KO cell line as well. This result suggests that not only does RER1 recognize misfolded PMP22 in post-ER compartments, but it may also recognize WT and other stable variants, returning them to the ER.

## Conclusions

In this work it was seen that elimination of N-glycosylation of PMP22 dramatically enhances the surface trafficking efficiency of both WT PMP22 and unstable forms of the protein such as the L16P CMTD mutant. We showed that the mechanisms of glycosylation differ for WT and disease mutant forms of PMP22, most likely due to an increased dwell time of less stable mutant forms of PMP22 at the Sec61 translocon. Indeed, WT and different mutants were seen to be N-glycosylated to differing degrees at the co-translational level (by OST-A) as opposed to the posttranslational level (via OST-B). As expected, CNX was seen to be an important contributor to this limiting role for N-glycosylation, although it was also seen that CNX plays glycosylation-independent roles. This work also clarified an important N-glycan independent role for RER1 in stringent Golgi-to-ER retrieval of PMP22 that has passed all other stages of ERQC. Our result also demonstrated, for the first time, that UGGT1 plays an important and glycosylation-dependent role in the ERQC of PMP22. Finally, this work provides a proteomic census of PMP22 interactive proteins that includes a readout for the differential levels of association with WT relative to both L16P and to glycosylation deficient N41Q PMP22. This work therefore both illuminates ERQC of PMP22 folding and trafficking and also provides information that will hopefully be useful in guiding future efforts to complete a map of the complex ERQC landscape experienced by PMP22 under conditions of both health and disease.

## Materials and Methods

For list of antibodies and guide RNA’s (gRNA’s) used in this study please see **Supplemental Table 2**.

### Cloning

Human cDNA for PMP22 was subcloned into a pIRES2 mammalian expression vector. To make PMP22 immunologically detectable we used QuikChange mutagenesis to insert a myc epitope into the second extracellular loop of PMP22 within the pIRES2 vector. QuikChange mutagenesis was also used to make the various point mutations used in this study. Plasmids were purified using a GenElute HP Plasmid MidiPrep Kit (Sigma-Aldrich). To generate plasmids for CRISPR/Cas9 KO cell lines, DNA encoding sgRNA’s were subcloned into pSpCas9-2A-Puro V2.0 (Addgene).

### Cell culture and transfection

HEK293 cells were obtained from the American Type Culture Center (ATCC), STT3A KO, STT3B KO, and MagT1/Tusc3 DKO HEK293 cells were a gift from Dr. Reid Gilmore. Cells were cultured in Dulbecco’s modified Eagle medium (DMEM) containing 10% fetal bovine serum (FBS) and 1% pen/strep at 37°C and 5% CO2. HEK293 cells were transiently transfected using the calcium phosphate method. 6 cm^2^ plates were transfected with 1.5 μg plasmid DNA. 10 cm^2^ plates were transfected with 5 μg plasmid DNA. Transfections were consistently >50% efficient.

### Single-cell trafficking assay

Single-cell flow cytometry was performed as outlined in REF. Briefly, ~36 hours post-transfection cells were trypsinized and prepared for FACS analysis using the Fix & Perm kit in accordance with the manufacturer instructions (Life Technologies, Grand Island NY). Cells were suspended in 100 μL of culture media, and a PE-labeled monoclonal anti-myc antibody was added to the solution to a final concentration of 0.75 μg/mL to immunostain surface PMP22. Cells were then incubated in the dark for 30 minutes at room temperature. 100 μL of fixation solution was then added to the media, and the cells were incubated in the dark for 15 minutes. The cells were then rinsed with 3 mL wash buffer (Phosphate buffered saline, PBS, containing 5% FBS and 0.1% NaN3) and pelleted via centrifugation twice. The cells were then suspended in 100 μL of permeabilization solution and incubated with an AlexaFluor647-labeled monoclonal anti-myc antibody at a final concentration of 0.75 μg/mL to label intracellular PMP22. Following 30 minutes of incubation in the dark, the cells were again rinsed with 3 mL wash buffer and pelleted via centrifugation twice. Cell were then resuspended in 300 μL wash buffer prior to FACS analysis.

Immunostained cells were analyzed with a FACS Canto II flow cytometer (BD Bioscience, San Jose, CA). Single cells were selected based their light scattering area and width profiles. 2500 transfected cells expressing PMP22 were analyzed from each sample by gating on GFP-positive cells (excited with a 488 nm laser, detected with 515–545 nm emission filter). The single-cell PE intensity (surface PMP22, excited with a 488 nm laser, detected with 564–606 nm emission filter) and Alexa Fluor 647 intensity (internal PMP22, excited with a 633 nm laser, detected with 650–670 nm emission filter) signals were corrected for nonspecific binding by subtracting the average intensities of untransfected, GFP-negative cells within each sample. To correct for the difference in the fluorescence intensity of the two antibodies, cells expressing WT PMP22 were stained with either the PE-labeled antibody or the Alexa Fluor 647-labeled antibody prior to FACS analysis, and the ratio of the average intensities of these cells was used to normalize the two signals. Single-cell trafficking efficiency values were then calculated from the ratio of the corrected PE signal of a given cell over the sum of its corrected Alexa Fluor 647 and PE signals. Average trafficking efficiency values calculated in this fashion were found to be similar to those determined by a comparison of the population-averaged intensities of intact (surface PMP22) and permeabilized (total PMP22) cells stained with the same concentration of the same fluorescently labeled antibody. Single-cell fluorescence intensity values below the background intensity were assigned an intensity of 0. Results were analyzed and visualized using FlowJo X software (Treestar Inc., Ashland, OR).

From titrations of both intact and permeable cells expressing WT PMP22 with fluorescently labeled antibodies, we found the average fluorescence intensity to be linearly dependent upon the antibody concentration. This confirms that fluorescence intensity values fall within the linear range of the detectors. Moreover, this ensures that the observed trafficking efficiency values are independent of the chosen antibody concentration. Compensation for spillover of the fluorescence signals between the channels utilized for the analysis as well as the gates for the selection of single cells, GFP-positive cells, and GFP-negative cells was initially set manually but was kept consistent for the collection of all data sets obtained thereafter.

### Co-IP and sample preparation for mass spectrometry

~36 hours post-transfection, confluent 10 cm^2^ plates were harvested via scraping in ice-cold PBS and pelleted via centrifugation. Cells were then lysed in lysis buffer (40 mM HEPES, 100 mM NaCl, 2 mM EDTA, 0.3% CHAPS, 10% glycerol, 1 mM PMSF, 1x HALT protease inhibitor, pH 7.8) for 20 minutes at 4°C with gentle rotation. Insoluble fractions were then removed via centrifugation and protein concentration determined via Bradford assay. 150 μg of total protein lysate was then mixed with 15 μL of anti-myc magnetic beads (Pierce) preequilibrated with lysis buffer and the volume brought up to 500 μL with TBS (25 mM Tris, 100 mM NaCl, pH 7.8). The mixture was then mixed with end-over-end rotation at 4°C for 90 minutes. Beads were then washed with 3x 250 μL volumes of TBS plus 0.25% Tween-20. Samples were then eluted with 2x 50 μL washes of elution buffer (50 mM Tris, 300 mM NaCl, 4% SDS, 2mM TCEP, 1 mM EDTA, pH 7.8) with beads vortexed and allowed to equilibrate with elution buffer for 15 min at 37°C for each wash.

Following elution, proteins were precipitated using chloroform/methanol extraction and protein pellets were allowed to air dry for 60 minutes at room temperature. Protein pellets were then resuspended in 50 μL 0.1% RapiGest SF (Waters, Milford, MA). Disulfide bonds were reduced with 1 mM TCEP and free sulfhydryl groups acetylated with 1 mM iodoacetamide and samples were incubated for 30 minutes at room temperature in the dark. Samples were then digested with 0.5 ug trypsin overnight at 37°C under 700 rpm shaking. Samples were then labeled using 6-plex tandem mass tags (TMT; Thermo Scientific, Waltham MA) according to the manufacturers protocol. TMT-labeled samples were then mixed and acidified with formic acid to a pH <2. Volume of the samples was then reduced to 1/6^th^ the initial volume on a speed-vac and then adjusted back to the original volume with Buffer A (H2O, 5% acetonitrile, 0.1% formic acid) to precipitate and remove RapiGest SF.

### Liquid chromatography – tandem mass spectrometry

MudPIT microcolumns were prepared as previously described ^53, 70^. Peptide samples were directly loaded onto the columns using a high-pressure chamber. Samples were then desalted for 30 minutes with buffer A (97% water, 2.9% acetonitrile, 0.1% formic acid v/v/v). LC-MS/MS analysis was performed using a Q-Exactive HF (Thermo Fisher) or Exploris480 (Thermo Fisher) mass spectrometer equipped with an Ultimate3000 RSLCnano system (Thermo Fisher). MudPIT experiments were performed with 10μL sequential injections of 0, 10, 30, 60, and 100% buffer C (500mM ammonium acetate in buffer A), followed by a final injection of 90% buffer C with 10% buffer B (99.9% acetonitrile, 0.1% formic acid v/v) and each step followed by a 130 minute gradient from 5% to 80% B with a flow rate of 300nL/minute when using the Q-Exactive HF and 500nL/minute when using the Exploris480 on a 20cm fused silica microcapillary column (ID 100 um) ending with a laser-pulled tip filled with Aqua C18, 3μm, 100 Å resin (Phenomenex). Electrospray ionization (ESI) was performed directly from the analytical column by applying a voltage of 2.0kV when using the Q-Exactive HF and 2.2kV when using the Exploris480 with an inlet capillary temperature of 275°C. Using the Q-Exactive HF, data-dependent acquisition of mass spectra was carried out by performing a full scan from 300-1800 m/z with a resolution of 60,000. The top 15 peaks for each full scan were fragmented by HCD using normalized collision energy of 38, 0.7 m/z isolation window, 120 ms maximum injection time, at a resolution of 15,000 scanned from 100 to 1800 m/z and dynamic exclusion set to 60s. Using the Exploris480, data-dependent acquisition of mass spectra was carried out by performing a full scan from 400-1600m/z at a resolution of 120,000. Top-speed data acquisition was used for acquiring MS/MS spectra using a cycle time of 3 seconds, with a normalized collision energy of 36, 0.4m/z isolation window, 120ms maximum injection time, at a resolution of 30,000 with the first m/z starting at 110. Peptide identification and TMT-based protein quantification was carried out using Proteome Discoverer 2.3 or 2.4. MS/MS spectra were extracted from Thermo Xcalibur .raw file format and searched using SEQUEST against a Uniprot human proteome database (released 03/2014 and containing 20337 entries). The database was curated to remove redundant protein and splice-isoforms, and supplemented with common biological MS contaminants. Searches were carried out using a decoy database of reversed peptide sequences and the following parameters: 10ppm peptide precursor tolerance, 0.02 Da fragment mass tolerance, minimum peptide length of 6 amino acids, trypsin cleavage with a maximum of two missed cleavages, dynamic methionine modification of 15.995 Da (oxidation), static cysteine modification of 57.0215 Da (carbamidomethylation), and static N-terminal and lysine modifications of 229.1629 Da (TMT sixplex). SEQUEST search results were filtered using Percolator to minimize the peptide false discovery rate to 1% and a minimum of two peptides per protein identification. TMT reporter ion intensities were quantified using the Reporter Ion Quantification processing node in Proteome Discoverer 2.3 or 2.4 and 576 summed for peptides belonging to the same protein.

### Interactome characterization and pathway enrichment analysis

To identify true interactors from non-specific background TMT intensities first underwent a log2 transformation and were then median normalized by dividing the intensity of each log2 TMT value by the median intensity in that channel to normalize across biological samples. TMT ratios were then calculated between by subtracting the log_2_ transformed and median corrected TMT intensity of the mock channels from the PMP22 channels. The mean log_2_ interaction differences were then calculated across multiple biological replicates. Significance of interaction differences were calculated using a paired, parametric, two-tailed t-test and multiple testing correction via an FDR estimation. Pathway enrichment analysis was performed using the DAVID bioinformatic server^55–56^.

To build the heat map shown in **Figure 4B**, peptide identifications were filtered to include those that were only identified in at least three biological replicates, had a log2 fold interaction greater than 0.2 and had a Q-Value <0.1 compared to mock samples. Identified proteins that passed this criteria were grouped via GO_term using the DAVID bioinformatic tool^55–56^. Manual curation of the GO_term analysis was then used to build the heatmap in a temporal manner based on literature precedent. GraphPad Prism was then used to generate the heat map to visualize the grouped interactions.

### Glycosylation mapping

For glycosylation mapping experiments cells were harvested and lysed as described above except for 60 minutes at 4° C. Following lysis, 5 μg samples of total lysate were treated with no enzyme, EndoH, or PNGase (New England Biolabs) according to the manufacturers protocol for 2 hours at 37°C. Samples were then analyzed via western blotting.

### Sucrose density fractionation

Cells were harvested and lysed as described in glycosylation mapping. Following lysis, 100 μg of cell lysate was layered on top of a 0-60% step-wise sucrose density gradient in 1 mL ultracentrifuge tube. Lysates were then centrifuged at 200,000 x g in a Beckman TLA-120.2 rotor for four hours at 4°C. Layers of the gradient were then sequentially removed in 100 μL fractions to be analyzed via western blotting.

### Generation of KO cell lines

CRISPR technology was utilized to generate genetic KO cells as previously described^71^. gRNAs were designed to target early exons of the CNX, Rer1, UGGT1, and LMAN1. gRNA oligomers (gRNA sequences shown in **Supplemental Table 2**) were annealed, phosphorylated, and ligated into digested pspCas9(BB)-2A-puro plasmid (plasmid no. 62988, Addgene, Cambridge, MA). HEK293 cells were suspended in 2 ml of DMEM supplemented with 10% FBS and plated in 6-well plates. The following day, 5 μg of each plasmid was combined with 10 μl Lipofectamine 2000 reagent (Invitrogen) in 1 ml Opti-MEM and incubated at room temperature for 30 min. Culture medium was replaced with the appropriate plasmid-lipofectamine solution, and cells were incubated at 37°C for 24 h. The medium was then replaced with fresh DMEM plus 10% FBS, and cells were allowed to recover for 24 h at 37°C prior to the addition of 0.75 μg/ml puromycin. Cells were incubated at 37°C for 48 h before replacing the medium. After recovering for ~1–2 weeks, cells were pelleted, resuspended in sorting buffer (PBS + 4% FBS), and strained to separate clumps of cells. Solutions were sorted by flow cytometry using a 5-laser BD LSRII with a 100 μm nozzle at the Vanderbilt Medical Center Flow Cytometry Core to isolate single cell cultures in 96-well plates for each cell line. Clones were incubated until they reached ~70% confluency and then passaged until enough cells could be harvested for KO validation via Western blotting.

### Statistics

Student’s t-test was used to compare mean values in trafficking experiments and glycosylation mapping experiments. Storey Q-values were used to compare identifications in the proteomics data. Pairwise Kolmogorov–Smirnov test was used to compare population distributions. Statistical analysies were performed in GraphPad Prism and Microsoft Excel.

## Data Availability

Proteomic data was deposited to the ProteomeXchange Consortium via the PRIDE partner repository with the identifier PXD023091. All other data are either presented in the article, are in the supporting information, or are available from the corresponding author upon request.

## Acknowledgements

We thank Arina Hadziselimovic for assistance with cloning and the staff of the Vanderbilt University Medical Center Flow Cytometry Shared Resource.

## Funding and Additional Information

This work was supported by National Institutes of Health Grants R01 NS095989 (to C. R. S.) and R35 GM133552 (L.P.). J. T. M. was supported by National Institutes of Health Fellowship F31 NS113494 and National Institutes of Health Training Grant T32 NS00749. M.T.W. was supported by National Institutes of Health Training Grant T32 GM065086, the Vanderbilt Institute of Chemical Biology Fellowship, and the National Science Foundation Graduate Research Fellowship Program. The Vanderbilt University Medical Center Flow Cytometry Shared Resource is supported by Vanderbilt Ingram Cancer Center Grant P30 CA68485 and Vanderbilt Digestive Disease Research Center Grant P30 DK058404. The content is solely the responsibility of the authors and does not necessarily represent the official views of the National Institutes of Health.

## Author Contributions

J.T.M. and C.R.S. conceptualized the work; J.T.M., M.T.W., L.P. and C.R.S. designed the methodology; J.T.M. and M.T.W. performed formal analysis; J.T.M M.T.W and D.R.H. performed investigation; L.P. and C.R.S. provided resources; J.T.M., M.T.W., L.P., and C.R.S. participated in data curation; J.T.M and C.R.S. wrote the original draft; all authors participated in reviewing and editing the manuscript; J.T.M. prepared the visualization; L.P. and C.R.S supervised the project; L.P. and C.R.S administered the project. J.T.M., M.T.W., L.P. and C.R.S acquired funding for the project.

## Conflict of Interest

The authors declare that they have no conflicts of interest with the contents of this article.

## Notes

http://proteomecentral.proteomexchange.org/cgi/GetDataset?ID=PXD023091

### Competing Interest Statement

The authors have declared no competing interest.

