## Supplemental Figures and Tables for "Glycosylation Limits Forward Trafficking of the Tetraspan Membrane Protein PMP22"

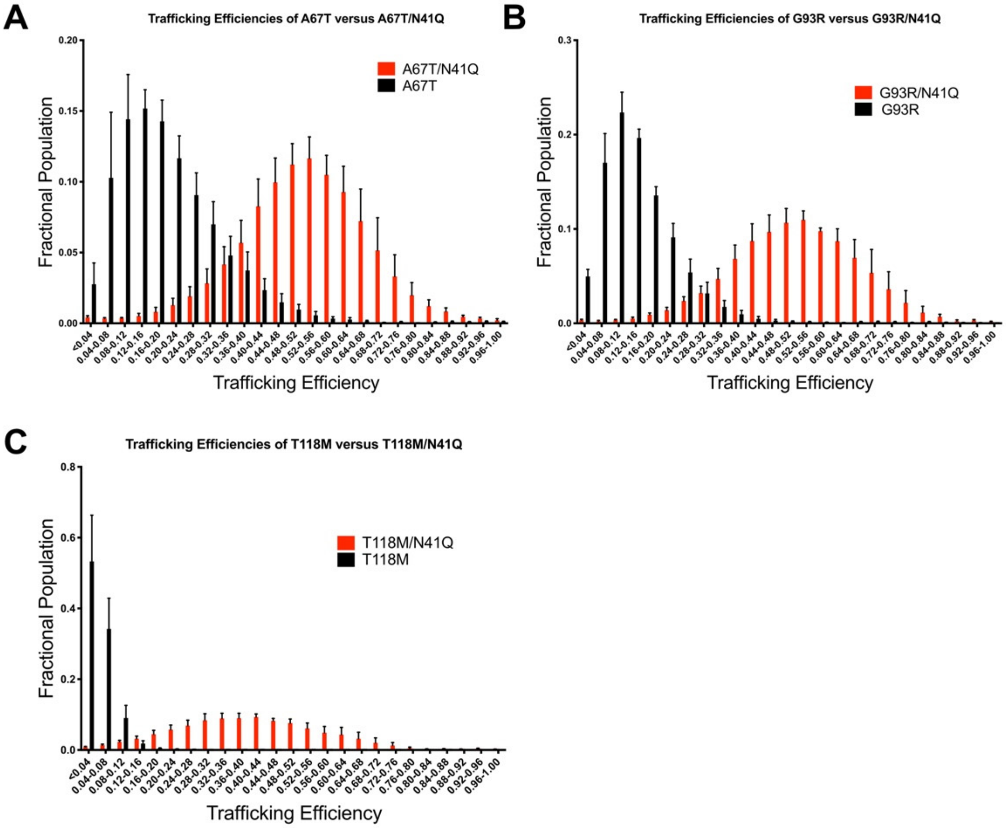


**Supplemental Figure 1**. Trafficking efficiencies of CMTD PMP22 variants and their glycosylation deficient parallels**.** Population distribution of PMP22 trafficking efficiencies measured in individual HEK293 cells for (**A**) A67T, (**B**) G93R, and (**C**) T118M PMP22. Values are shown for glycosylated (black) and non-glycosylated (red) variants. Measurements were obtained from 5 biological replicates with 2500 cells measured per replicate. Error bars represent standard deviations (SD) of the replicates.

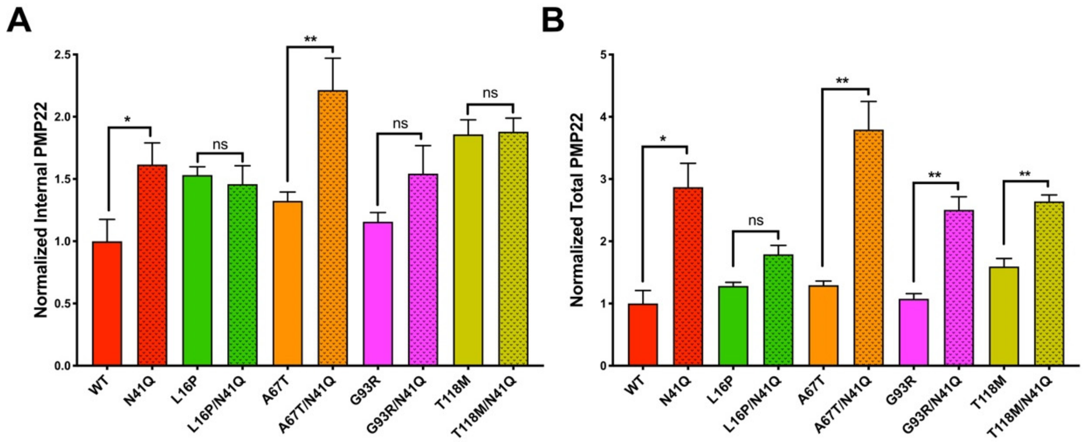


**Supplemental Figure 2**. Normalized internal and total PMP22 concentrations. Normalized **(A)** internal and **(B)** total expression of PMP22 variants and their glycosylation deficient partner. Values were obtained from 5 biological replicates with 2500 cells measured per replicate. All values were normalized to WT PMP22 cell surface expression data collected in paired biological replicates. Error bars represent SD of the replicates. Student's t-test was used for statistical analysis. ns=not significant, *=$p<0.05$, **=$p<0.01$.

A

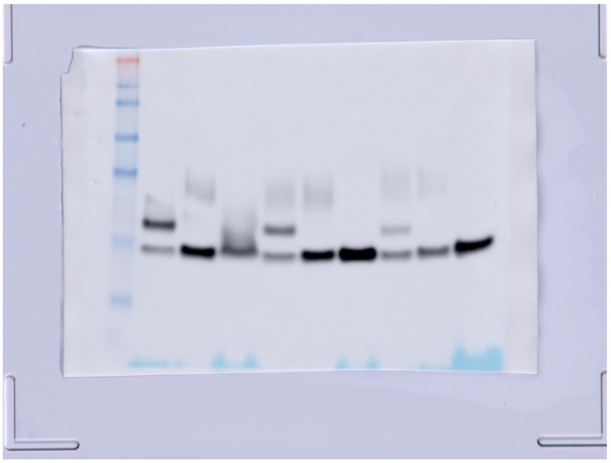

B

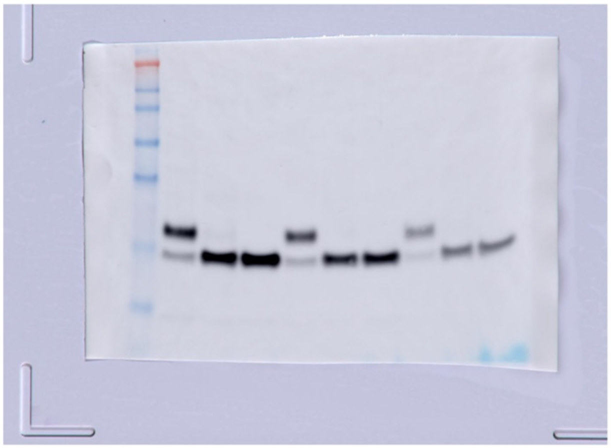

**Supplemental Figure 3**. Uncut Western blots from **(A) Figure 2A** and **(B) Figure 2B.**

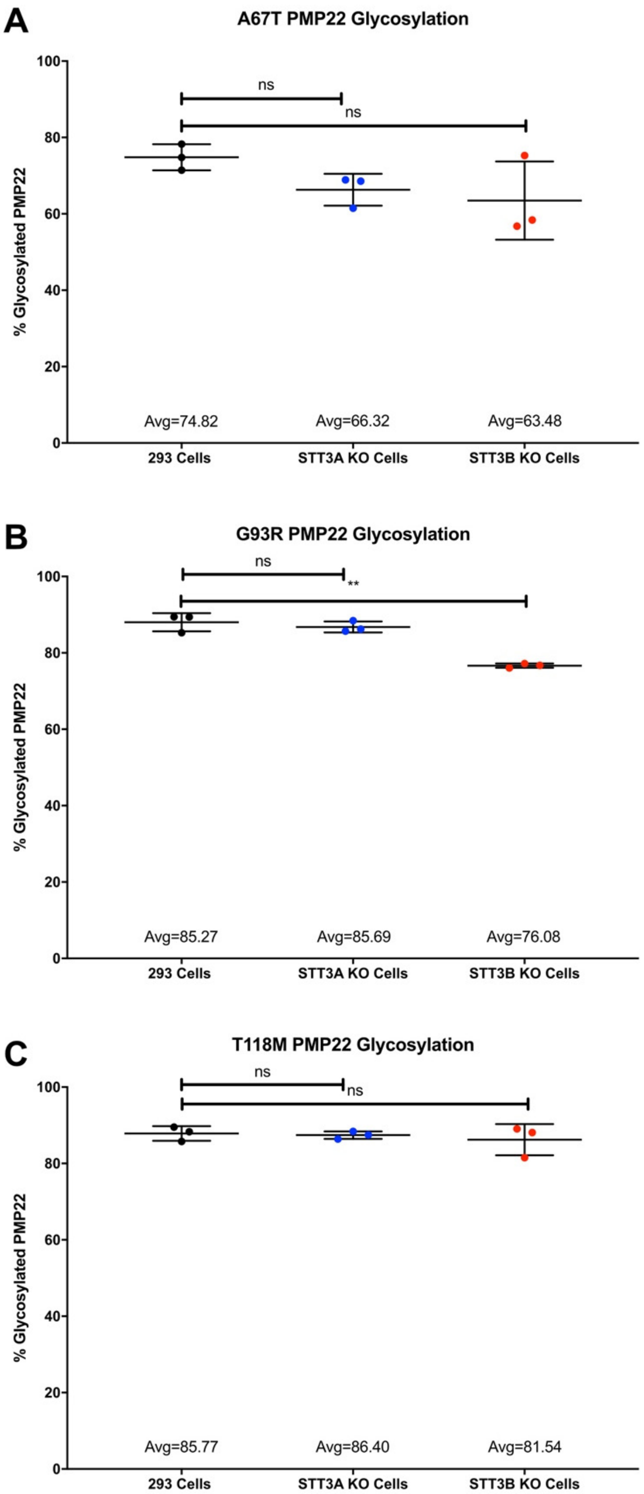


**Supplemental Figure 4.** Glycosylation mapping for CMTD PMP22 variants. Quantified levels of glycosylated **(A)** A67T, **(B)** G93R, and **(C)** T118M PMP22 from 3 independent biological replicates.

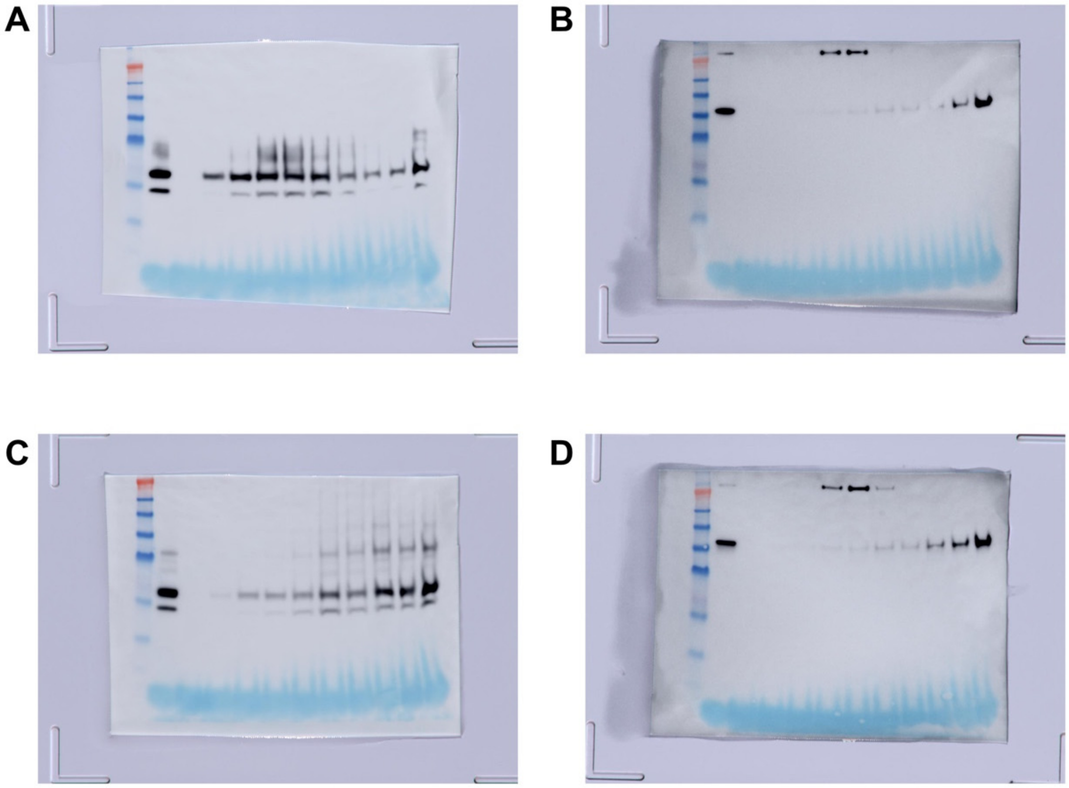


**Supplemental Figure 5.** Uncut Western blots from **(A-B) Figure 2C** and **(C-D) Figure 2D**.

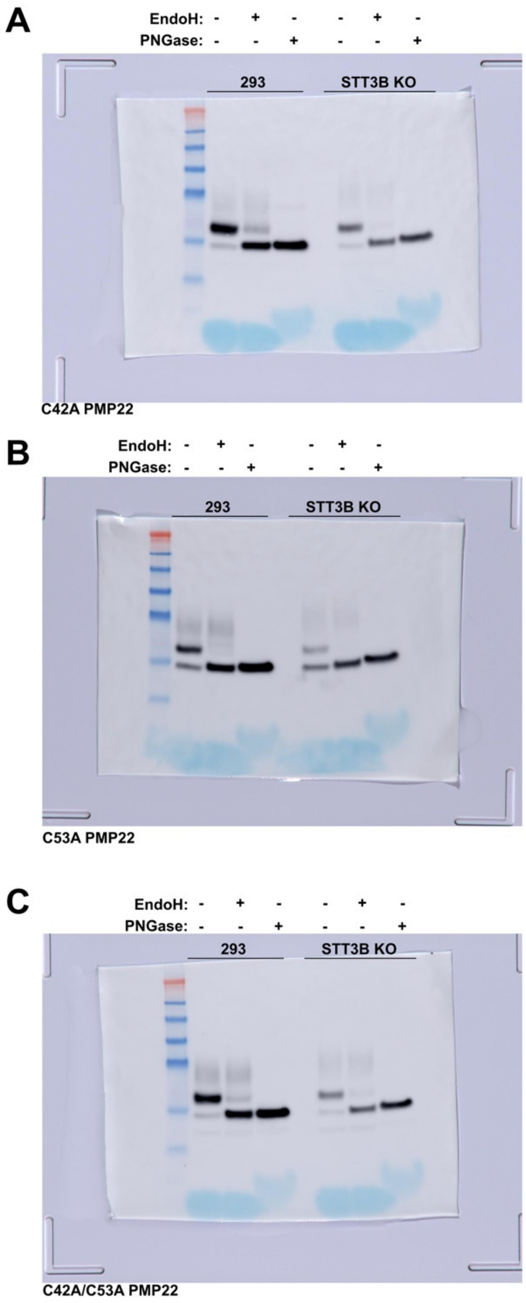


**Supplemental Figure 6**. Western blots showing the levels of glycosylation of **(A)** C42A, **(B)** C53A, and **(C)** C42A/C53A PMP22 in HEK293 or STT3B KO HEK293 cells.

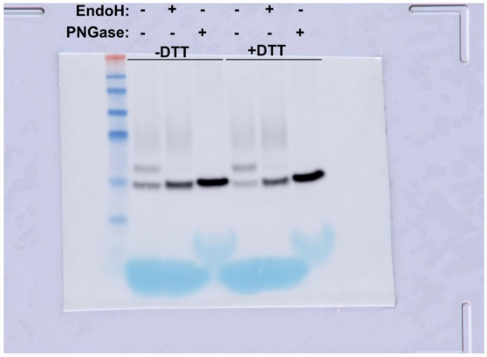


**Supplemental Figure 7.** Western blot showing the level of WT PMP22 glycosylation in STT3B KO HEK293 cells that have been pretreated for 2 hours with or without 2 mM DTT.

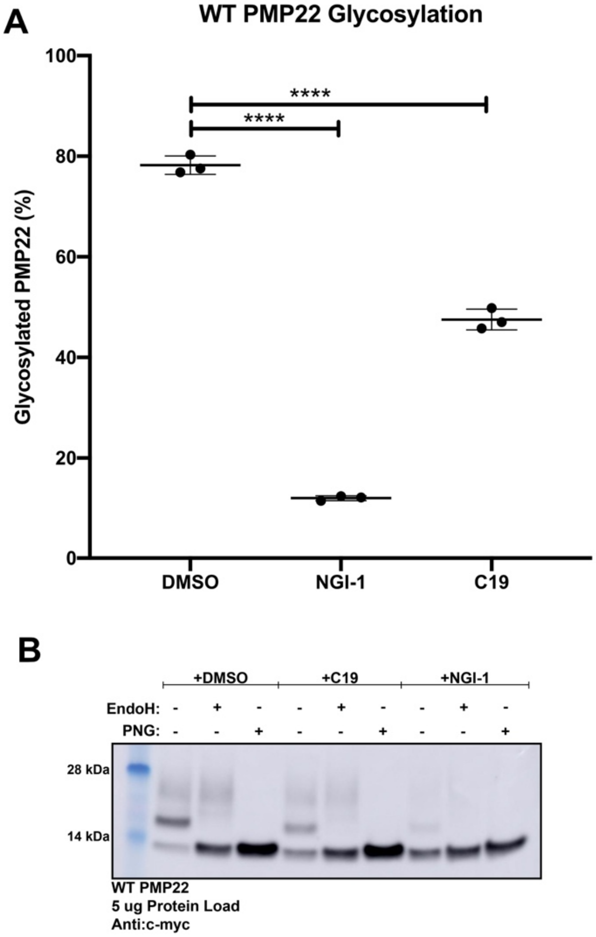


**Supplemental Figure 8**. WT PMP22 glycosylation with OST inhibitors. **(A)** Quantification of the levels of WT PMP22 glycosylation from three biological replicates for lysates obtained from cells that had been treated for 24 hours with either DMSO, 10 μM NGI-1, or 10 μM C19. Error bars represent standard deviation. **(B)** Representative Western blot showing the levels of WT PMP22 glycosylation from lysates obtained from cells that had been treated for 24 hours with either DMSO, 10 μM NGI-1, or 10 μM C19. ****=p<0.0001 using student's t-test.

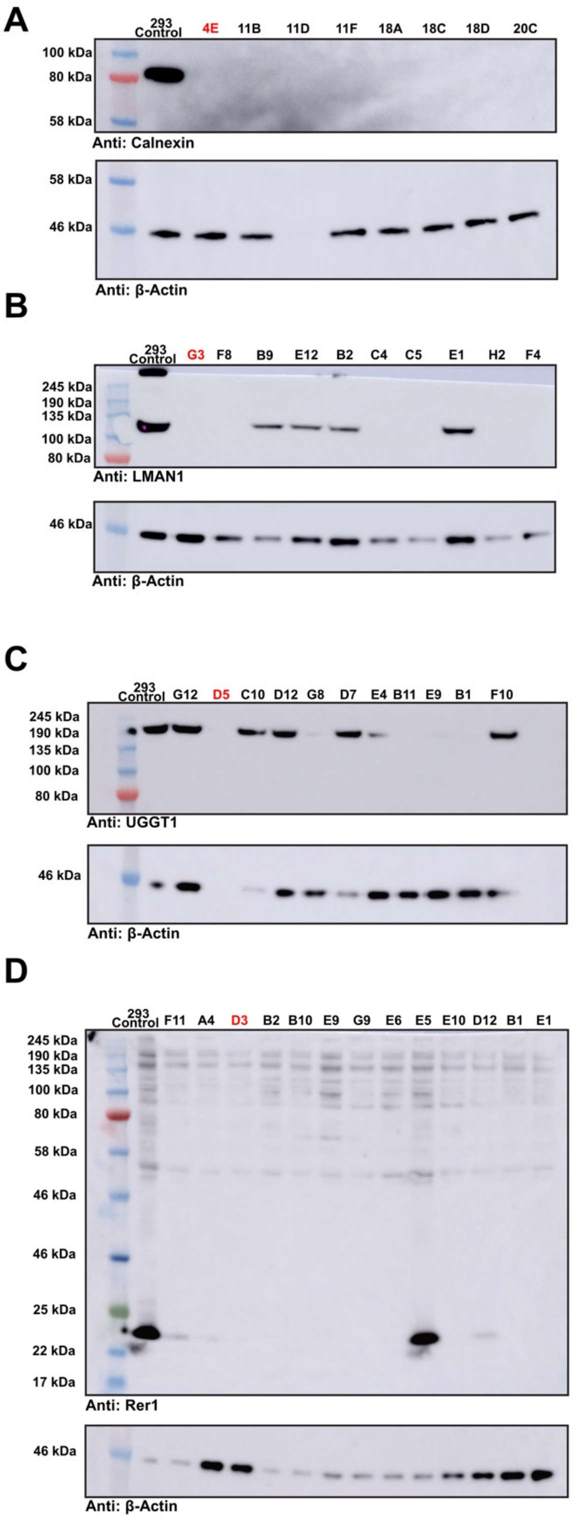


**Supplemental Figure 9**. Western blots of CRISPR/Cas9 clonal KO cell lines for **(A)** calnexin (CXN), **(B)** LMAN1, **(C)** UGGT1, and **(D)** RER1. β-Actin was used as a loading control for all samples. Clones selected for trafficking studies are highlighted in red.

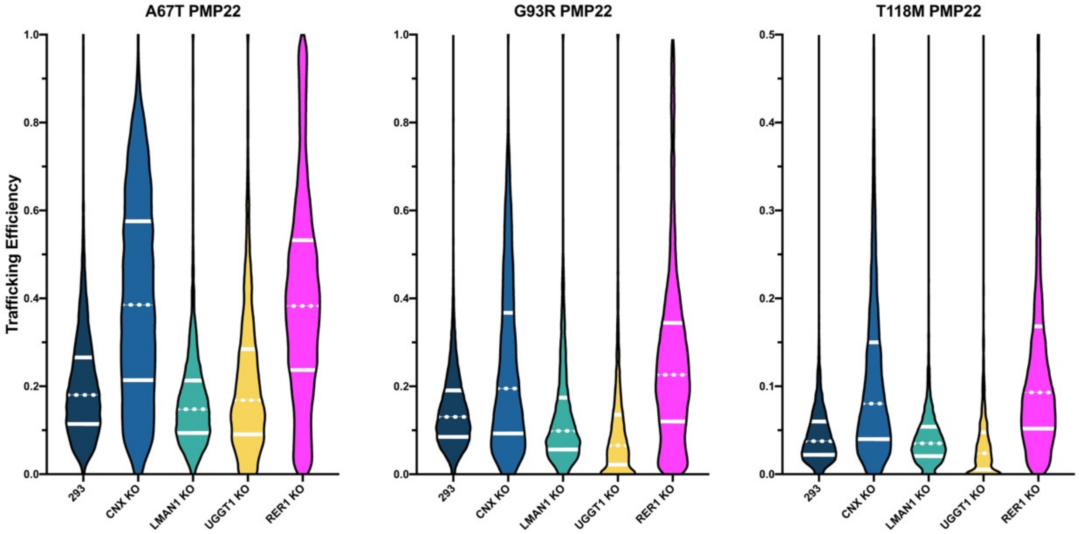


**Supplemental Figure 10**. PMP22 trafficking efficiencies in CRISPR/Cas9 KO cells of potential ERQC interactors. CRISPR/Cas9 was used to generated KO cells of potential proteins involved in mediating trafficking effects for PMP22. Violin plots showing population distributions of A67T, G93R and T118M PMP22 trafficking efficiencies from three biological replicates are shown. Data was collected in HEK293 cells (navy), CNX KO HEK293 cells (blue), LMAN1 KO HEK293 cells (green), UGGT1 KO HEK293 cells (yellow) and RER1 HEK293 KO cells (pink). White lines in the population distributions separate the data into quartiles.

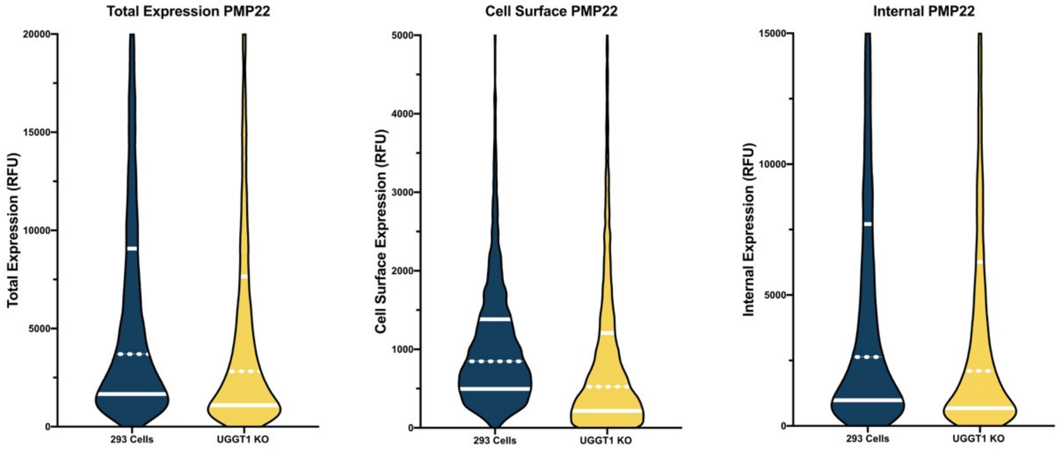


**Supplemental Figure 11**. PMP22 total, cell surface, and internal expression in HEK293 or UGGT1 KO cells. Violin plots showing population distributions of WT PMP22 total, cell surface, or internal expression levels from three biological replicates are shown. Data was collected in HEK293 cells (navy), or UGGT1 KO HEK293 cells (yellow). White lines in the population distributions separate the data into quartiles, with the dotted line representing the median.

```
P16646      PMP22_MOUSE        1   MLLLLLGILFLHIAVLVLLFVSTIVSQWLVGNGHTTDLWQNCTTSALGAVQHCYSSSVSE   60
P25094      PMP22_RAT          1   MLLLLLGILFLHIAVLVLLFVSTIVSQWLVGNGHRTDLWQNCTTSALGAVQHCYSSSVSE   60
Q01453      PMP22_HUMAN        1   MLLLLLSIIVLHVAVLVLLFVSTIVSQWIVGNGHATDLWQNCSTSSSGNVHHCFSSSPNE   60
Q9TQZ3      PMP22_BOVIN        1   MLLLLGIIVLHVAVLVLLFVSTIVSQWMVGNGHATDLWQNCSTSLMGSVQHCFSSSANE    60
Q6WL85      PMP22_HORSE        1   MLLLLLGIIVLHVAVLVLLFVATIVSQWIVGNGHATDLWQNCSTT-SGNVQHCLSSSANE   59
U3K4I0      U3K4I0_FICAL       1   MLLLLLGIIVLHVTVLVLLFVSTIVSQWLVNGEHRADLWQNCTS---GSSFHCLASSSNE   57
A0A452E2E7  A0A452E2E7_CAPHI  1   MLLLLLGIIVLHVAVLVLLFVSTIVSQWMVGNGHATDLWQNCSTSSMGSVQHCFSSSANE   60
Q6DCV9      Q6DCV9_XENLA       1   MLLLLLGIIVLHIAILVLLFVSTIVSSWLVGNGYSADLWQNCSGPATG-TWQCLTSSSNE   59
                                   ******.*:.**:::******:****.*:*.. : :******:    *   :* :** .*
```

**Supplemental Figure 12**. Sequence alignment of PMP22 from eight different species. Highlighted region shows the glycosylation motif.
